# Effects of selfing on the evolution of sexual reproduction

**DOI:** 10.1101/2023.03.07.531539

**Authors:** Kuangyi Xu

## Abstract

Plants exhibit diverse breeding systems, with populations capable of outcrossing, selfing, and/or asexual reproduction. However, interactions between the three reproductive pathways remain not fully clear. Sexual reproduction introduces segregation and recombination, but incurs several costs. Selfing can affect the relative costs and benefits of sexual vs. asexual reproduction. Building population genetic models, I explore how selfing affects the evolution of a sexual reproduction rate modifier via (1) indirect selection due to segregation, (2) indirect selection from changes in recombination rates, and (3) selection from the cost of meiosis and mate limitation. The dominant selective force mediating the evolution of sex is found to vary with the rate of sexual reproduction and selfing, but selective force (1) and (3) are generally stronger than selective force (2). A modifier enhancing sexual reproduction tends to be favored by indirect selection generated by partially recessive, small-effect deleterious mutations, while hindered by highly recessive lethal mutations. Overall, evolution towards higher sexual reproduction is hindered at low sexual reproduction rates and intermediate selfing rates, but favored under high selfing rates. The results suggest that asexual reproduction may precede the evolution of selfing, and offer insights into the evolution of mechanisms reducing geitonogamy in partially clonal populations.

## Introduction

Plants can reproduce through outcrossing, self-fertilization, and asexual reproduction. Many plant populations reproduce through a combination of the three modes. As most plants are hermaphroditic, selfing occurs in more than 50% of plant species (Jarne and Charlesworth 1993, Goodwillie et al. 2005). Asexual reproduction is also prevalent in plants and occurs mainly in two ways: vegetative reproduction and apomixis. Vegetative reproduction is very common in plants (Gustafsson, 1947, Abrahamson 1980, Silvertown 1987), which can be realized through budding from organs such as roots, stems, leaves, and inflorescences (Eckert 1999, Silvertown 2014, Barrett 2015). Apomixis refers to asexual seed production (Asker and Jerling 1992, Richards 2003, Whitton et al. 2008, Hojsgaard and Hörandl 2019), but it is much rarer compared to vegetative reproduction (Whitton et al. 2008). Also, unlike vegetative reproduction, apomixis seems to be unlikely to coexist with sexual reproduction within the same population (Bengtsson and Ceplitis 2000, Nakayama et al. 2002, Whitton et al. 2008).

To understand the diversity of breeding systems in plants, it is important to know how asexual reproduction, outcrossing, and selfing interact to affect the evolution of each other. Empirical studies show that apomictic populations often derived from self-incompatible lineages (Hörandl 2010, Hojsgaard and Hörandl 2019), and many facultatively asexual reproducing species are outcrossing (Ellstrand and Roose 1987). Nevertheless, partially clonal populations can still exhibit a wide range of selfing rates (Honnay and Jacquemyn 2008, Vallejo-Marín et al. 2010), with some being highly self-fertilizing (e.g., *Iris versicolor*, Kron et al. 1993). Therefore, how self-fertilization and asexual reproduction interact with each other is still not well understood. This is partly because studies on the evolution of selfing often assume complete sexual reproduction, while studies on the evolution of sex usually consider random mating.

The evolution of sexual reproduction has attracted long-lasting interest in the field of evolutionary biology. Although sexual reproduction is prevalent, it bears several costs compared to asexual reproduction (reviewed in Lehtonen et al. 2012). Table 1 summarizes several key selective forces on sex and their causes. One of the major costs of sex is the “cost of males”. In a sexually reproducing population, females allocate resources to produce males in separate-sex populations (or the male function in hermaphrodites), but male gametes contribute few resources to the offspring zygote. Also, mate search and choice require extra resources and energy of females (Smith and Maynard-Smith 1978, Lehtonen et al. 2012, Meirmans et al. 2012). Such costs have been observed in various taxa (Michaels and Bazzaz 1986, Innes et al. 2000, Kumpulainen et al. 2004, Wolinska and Lively 2008). However, the cost of males can be compensated by benefits provided by males such as parental care, or increased access to resources for offspring production (Lehtonen et al. 2012). Another cost of sex is the “cost of meiosis” (Williams 1975) or the cost of “genome dilution” (Lewis 1987). Specifically, in a hermaphroditic population, asexually reproducing individuals can transmit more alleles to the next generation than a sexually reproducers, by siring female gametes of both theirs own and other sexually reproducers. This cost of sex is weaker if increased allocation to asexual reproduction reduces the amount of male gametes that sire other individuals. The cost of males can occur for both within-population and between-population selection on sex, and may apply to both hermaphroditic and separate-sex populations. In contrast, the cost of meiosis only occurs for within-population selection on sex, and only applies to hermaphroditic populations. For intuition, in separate-sex populations, asexually reproducing females do not possess the male function to fertilize sexually reproducing females, and thus are genetically isolated from sexual individuals (see Box 2 and 3 in Lehtonen et al. (2012) for detailed explanations).

**Table 1:** A summary of selective forces on a sex modifier and the effects of selfing on each force.

| Selective forces on a sex modifier | Causes of the selective force | How selfing affects the selective force |  | Related figures in this paper |
| --- | --- | --- | --- | --- |
| Indirect selection due to interactions between alleles within a selected locus (intra-locus interaction) | Changes in the rate of segregation | Alter the fitness of sexually reproduced offspring relative to asexually reproduced offspring | 1. An excess of homozygosity at a locus<br>2. Generate positive association between homozygotes at different loci.<br>3. Alter genetic association between the sex modifier locus and fitness loci | Fig. 1, 2, 3 |
| Indirect selection due to interactions between selected loci (inter-locus interactions) | Changes of the effective recombination rate across the genome | Reduce the effective recombination rate in the genome |  | Fig. 4 |
| Cost of males | Resources cost to produce males or the male function | Reduce resource allocation to the male function |  | N/A |
| Cost of meiosis | Sexual individuals transmit only half of the genome to the offspring | Reduce the cost of meiosis as a mode of uniparental reproduction |  | Fig. 6(b) |
| Mate limitation | Outcrossing requires availability of gametes from other individuals | Reduce the strength of mate limitation by providing reproductive assurance |  | Figs. 6(a), 6(b) |

Despite the costs reviewed above, sexual reproduction is common in eukaryotes (Bell 1982), so numerous studies have tried to identify the selective advantages of sex. Sexual reproduction differs from asexual reproduction by two genetic characteristics. First, sexual reproduction allows the possibility of recombination. A sex modifier can thus be indirectly selected due to a change in the effective recombination rate under interactions between selected loci, hereafter referred to as selection caused by inter-locus interactions. Previous studies show that an allele increasing the recombination rate can be indirectly selected for by breaking negative linkage disequilibria between loci, thereby enhancing the efficiency of selection. This outcome may arise from epistatic interactions between loci (Charlesworth et al. 1979, Barton 1995, Roze and Lenormand 2005, Roze and Stetsenko, Roze 2022) or the interplay between selection and drift in finite population size, known as the Hill-Robertson effect (Hill and Robertson 1966, Otto and Barton 2001, Keightley and Otto 2006, Roze and Barton 2006, Roze 2021).

Secondly, in a diploid population, sexual reproduction entails segregation. Consequently, sex can be favored under interactions between alleles within a locus, referred to as selection caused by intra-locus interactions. Specifically, consider a fitness locus with allele *A* and *a*, where *a* is deleterious. Asexual reproduction will generate an excess of heterozygotes than that under Hardy-Weinberg equilibrium, when allele *a* is recessive enough so that the heterozygote *Aa* is on average fitter than homozygotes (i.e., 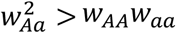) (Chasnov 2000, Agrawal and Chasnov 2001), or when the population size is finite (Roze and Michod 2010). This intra-locus association reduces genetic variance and hinders selection against the deleterious allele *a*. A modifier allele that increases the rate of sexual reproduction can gain a long-term advantage by breaking this association through segregation. This is because it augments genetic variance and thus enhancing mean fitness at equilibrium, as selection becomes more efficient in eliminating the deleterious allele (Otto 2003). However, the modifier will face a short-term disadvantage since it immediately produces more homozygous offspring, which has a lower average fitness than heterozygotes.

Regarding the multiple selective forces on the evolution of sex reviewed above, selfing can have various effects on the relative costs and advantages of sexual vs. asexual reproduction, as summarized in Table 1. First, selfing is a mode of uniparental reproduction, thus reducing the cost of meiosis during sexual reproduction (Charlesworth 1980). Additionally, selfing serves as an alternative method to asexual reproduction for providing reproductive assurance under mate limitation, where outcrossing is disfavored (Busch and Delph 2012, Eckert et al. 2016).

Also, selfing influences indirect selection on sex through changes in the recombination rate caused by inter-locus interactions. Selfing generates positive associations among homozygotes at different loci, referred to as identity disequilibrium (Vitalis and Couvet 2001). This effect can lead to selection against a modifier increasing the recombination rate in the absence of epistasis. However, it can generate indirect selection for a higher recombination rate under negative epistatic interactions between deleterious mutations (Charlesworth et al. 1979, Roze and Lenormand 2005, Stetsenko and Roze 2022). Indirect selection for a higher recombination rate can also be caused by the Hill-Robertson effect, which tends to be stronger under selfing for two reasons (Roze 2021, Stetsenko and Roze 2022). First, selfing reduces the effective recombination rate across the genome due to increased homozygosity. Second, selfing reduces the effective population size by increasing the variance in allele frequency changes (Caballero and Hill 1992) and strengthening background selection (selection against linked deleterious mutations; Stephan 2010, Charlesworth 2013). Nevertheless, previous studies on the evolution of the recombination rate often assume complete sexual reproduction, and the effects of selfing under the presence of asexual reproduction are not fully clear.

Furthermore, selfing will affect indirect selection on sex through segregation caused by intra-locus interactions. One reason is that selfing can alter genetic associations between the sex modifier locus and fitness locus (Otto 2003). Another factor is that selfing may change the relative fitness of sexually versus asexually reproduced offspring due to increased homozygosity (Muirhead and Lande 1997), thereby affecting the short-term (dis)advantage of a sex modifier. This effect is akin to inbreeding depression, where selfed offspring tends to have lower fitness than outcrossed offspring because selfing exposes partially recessive deleterious mutations to homozygotes (Charlesworth and Willis 2009). A higher selfing rate can lower inbreeding depression by purging deleterious mutations (Roze 2015). However, it is not clear whether selfing will decrease or increase the relative fitness of sexually reproduced compared to asexually reproduced offspring.

As for the overall effects of selfing on indirect selection caused by intra-locus interactions, Otto (2003) showed that gametophytic selfing (a mode of selfing in some mosses and ferns that involves the union of two gametes from the same haploid gametophyte; Soltis and Soltis 1992) can qualitatively alter the genetic conditions for a sex modifier to invade compared to those under outcrossing. However, in most plants and some animal groups, self-fertilization occurs through the union of two gametes from the same diploid sporophyte (referred to as sporophytic selfing, Jarne and Charlesworth 1993). Since gametes from the same haploid parent are identical, gametophytic selfing always produces homozygotes. In contrast, sporophytic selfing can still generate heterozygous offspring if the parent is heterozygous. Therefore, the effects of sporophytic selfing on indirect selection on sex through segregation are not fully clear. Uyenoyama and Bengtsson (1989) showed that sporophytic selfing can promote the evolution of sex, assuming the deleterious mutation is completely lethal. However, since the lethal allele only exists in the heterozygous state in their recursion equations, whether the results obtained in this special case hold for more general genetic conditions remains to be investigated.

Due to the complicated effects of selfing on the evolution of sex, we still lack have a comprehensive understanding of the relative importance of these forces, and the overall effects of selfing on the evolution of sex. Answering these questions requires quantifying the strength of each selective force. Nevertheless, previous studies that compare the strength of different selective forces on sex assume random mating (e.g., Roze and Michod 2010, Roze 2014). Therefore, in this paper, I use population genetic models and individual-based simulations to investigate the effects of selfing on the evolution of a sex modifier through: (1) indirect selection through segregation caused by intra-locus interactions (2) indirect selection caused by inter-locus interactions due to changes in the effective recombination rate, and (3) selection due to the cost of meiosis and mate limitation.

I find that for indirect selection caused by intra-locus interactions, under sporophytic selfing, a modifier increasing the rate of sexual reproduction is favored either when the deleterious allele is recessive and selection is weak, or when the deleterious mutation is dominant enough and selection is strong. This is qualitatively similar to results found under gametophytic selfing (Otto 2003). Moreover, since deleterious are often recessive, a higher selfing rate lowers the fitness of sexually reproduced offspring relative to asexually reproduced offspring, generating a short-term disadvantage to a modifier increasing the rate of sex due to segregation. Using empirically estimated genetic parameters of deleterious mutations, I show that indirect selection caused by intra-locus interactions has comparable strength to selection due to the cost of meiosis and mate limitation. Both selective forces are stronger than indirect selection caused by inter-locus interactions, unless the selfing rate is high. Overall, the evolution towards a higher sexual reproduction rate tends to be disfavored under an intermediate selfing rate and a low sexual reproduction rate, but is favored when the selfing rate is high. Given that clonality is often associated with outcrossing in plant populations, the current results suggest that asexual reproduction may evolve prior to the evolution of selfing in plant populations.

## Methods

This section consists of three subsections. The first and second subsections present a population genetic model for selection on a sex modifier caused by intra-locus interactions and inter-locus interactions, respectively. The third subsection describes the algorithm of individual-based simulations. The biological meanings of key symbols used in the model are summarized in Table 2. The Mathematica Notebook for all population genetic models and related analyses, as well as the C++ code for individual-based simulations, are available on Zenodo at https://doi.org/10.5281/zenodo.10479940.

**Table 2.**
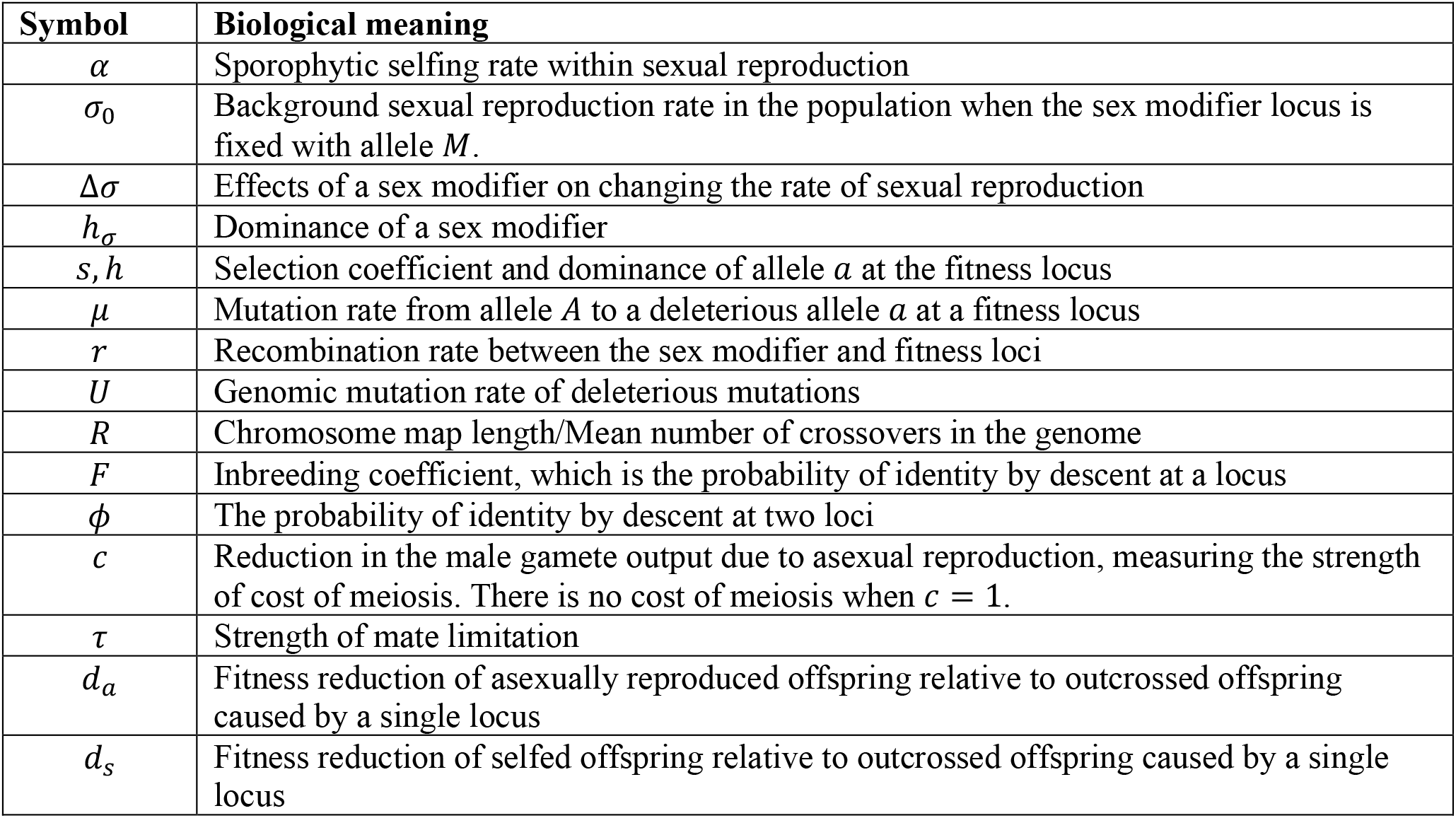
Biological meaning of key symbols used in the model.

### Indirect selection through segregation caused by intra-locus interactions

I consider a diploid population with an infinitely large population size. Each generation begins with viability selection, followed by mutations and reproduction. During reproduction, there may be selection due to the cost of meiosis and mate limitation. I consider a sex modifier locus **M** controlling the proportion of sexually reproduced offspring, and a fitness locus **A** governing individual viability, with a recombination rate *r* between them.

The sex modifier locus **M** has two alleles *M* and *m*. The rate of sexual reproduction of genotype *MM, Mm* and *mm* is *σ*_0_, *σ*_1_ = *σ*_0_ + *h*_*σ*_Δ*σ, σ*_2_ = *σ*_0_ + Δ*σ*, respectively. The model focuses on the invasion of allele *m* in a population previously fixed with allele *M*, so I refer to *σ*_0_ as the background sexual reproduction rate. During sexual reproduction, the sporophytic selfing rate is denoted by *α*. The fitness locus **A** has two alleles *A* and *a*, where allele *a* is deterious. The fitness of *AA, Aa* and *aa* are 1, 1 – h*s*, and 1 – *s*, respectively, where *s* and *h* are the selection coefficient and dominance of allele *a*. Allele *A* mutates to *a* with a rate *μ*, and I assume there is no backward mutation from *a* to *A*. I denote the genotype frequency at the beginning of each generation by *x*_*ij*_, where *i* ≤ *j* and *i, j* = 1,2,3,4 (1 for *MA*, 2 for *Ma*, 3 for *mA*, and 4 for *ma*).

Denote the genotype frequency after viability selection as 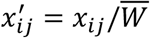, where 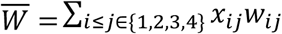 is the mean fitness, and *w*_*ij*_ is the fitness of genotype *x*_*ij*_. The genotype frequencies after mutation and selection, denoted by 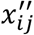, are given by

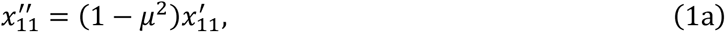

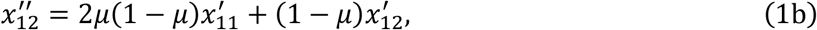

etc., and the frequency of other genotypes can be expressed in a similar way.

After selection and mutation, reproduction occurs. I denote the genotype frequency within asexually reproduced, selfed and outcrossed offspring by 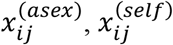 and 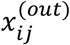, respectively. The genotype frequency within asexually reproduced offspring is 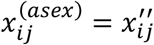. Under sporophytic selfing, the genotype frequency within selfed offspring can be calculated as

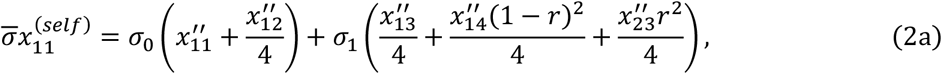

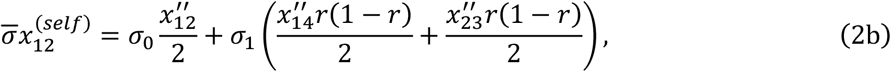

etc. Here, 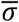 is the average sexual reproduction rate, given by

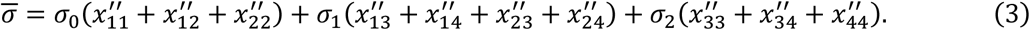

The frequency of other genotypes 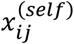 can be obtained in a similar way.

For outcrossed offspring, I denote the haplotype frequency within female gametes (i.e., ovules) and male gametes (i.e., pollen) by *y*_*i*_ and *z*_*i*_, respectively, which are calculated as

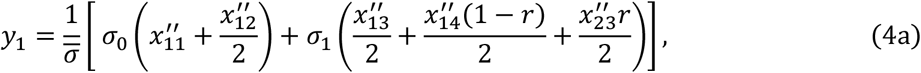

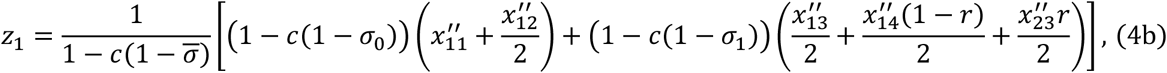

respectively. The frequency of other haplotypes can be calculated in a similar way. In equation (4a), I assume an increased rate of asexual reproduction causes a proportional reduction in female gamete production. In equation (4b), the parameter *c* measures the reduction in male gamete production due to an increased rate of asexual reproduction. When *c* = 1, an increased rate of asexual reproduction causes a proportional reduction in male gamete production (thus *y*_*i*_ = *z*_*i*_), so the sex modifier is neutral when there is no fitness locus. When *c* < 1, sex suffers the cost of meiosis since asexual reproduction can transmit extra male gametes. I assume pollination is random, so the genotype frequency within outcrossed offspring is

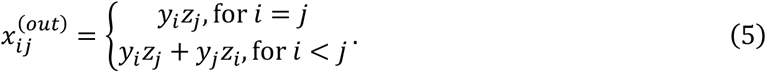

The reproductive success of outcrossing, selfing and asexual reproduction, as well as the fitness of offspring produced from the three reproductive modes may differ due to factors other than the focal fitness locus (e.g., mate limitation, different seed provisioning). To account for these effects, I use *w*_*asex*_ and *w*_*self*_ to represent the reproductive success and the offspring fitness of asexual reproduction and selfing relative to outcrossing. The genotype frequency in the next generation is thus given by

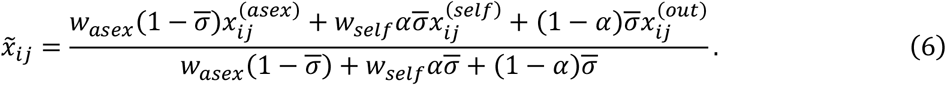

To investigate whether a sex modifier allele *m* can invade into a population previously fixed with allele *M*, I numerically calculate the leading eigenvalue *λ* of the Jacobian matrix given by equation (6) (a complicated analytical expression of *λ* is available in Section 3.3 of the Mathematica Notebook). *m* can invade when *λ* > 1. I also track the genetic associations within loci and between the modifier and fitness locus after the introduction of a modifier allele *m* in the population, to see how genetic associations affect the capacity for a sex modifier to invade (the calculation of genetic associations is available in Section 1.3 of the Mathematica Notebook).

To estimate the short-term (dis)advantage of a sex modifier due to fitness differences between sexually and asexually reproduced offspring caused by the focal fitness locus, I calculate the fitness of asexually reproduced offspring and selfed offspring relative to outcrossed offspring contributed by the focal locus. To do this, I first solve the genotypic frequencies at the fitness locus under mutation-selection balance based on equation (6). The mean fitness of asexually reproduced offspring is merely equal to the mean fitness of the population 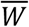. The mean fitness of selfed and outcrossed offspring can be calculated as

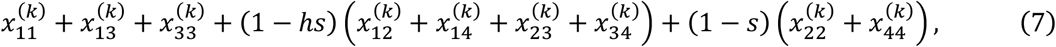

where *k* = *self* or *out*, and 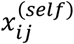 and 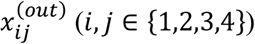 is given by equations (2) and (5), respectively.

### Indirect selection caused by inter-locus interactions

Indirect selection on sex through changes in the recombination rate caused by inter-locus interactions can occur due to two causes: deterministic interactions between loci and the Hill-Robertson effect in a finite population. I present models for estimating the strength of selection caused by deleterious mutations in the genome under each cause, respectively.

#### Deterministic interactions between loci

To estimate the strength of indirect selection caused by deterministic interactions between deleterious mutations in an infinitely large population, I modify the model described in Stetsenko and Roze (2022). Briefly, I consider two chromosomes with map length *R*, and assume the loci where deleterious mutations occur are uniformly distributed along the chromosome with a genomic mutation rate *U*. A sex modifier locus *i* is located at the midpoint of the chromosome. I assume the recombination rate between locus *j* and *k* is 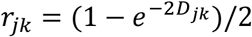 (Haldane 1919), where *D*_*jk*_ is the genetic distance between locus *j* and *k*. For a sex modifier that changes the rate of sexual reproduction by Δ*σ*, it effectively changes the map length by Δ*R* = *R*Δ*σ*. The modifier thus effectively changes the recombination rate between locus *j* and *k* by Δ*r*_*jk*_ = *r*_*jk*_Δ*R*/*R*.

I denote the strength of selection on the modifier locus *i* contributed by fitness locus *j* and *k* as *s*_*jk*_(*D*_*ij*_, *D*_*jk*_, *D*_*ik*_), where *D*_*ij*_, *D*_*jk*_ and *D*_*ik*_ are genetic distance between pairwise loci. A complicated expression of *s*_*jk*_ is available in Section 4.1 of the Mathematica Notebook. Briefly, *s*_*ij*_ is obtained using equation (6) in Stetsenko and Roze (2022), by replacing the probability of identity by descent at one locus *F* (inbreeding coefficient) and at two loci *ϕ* with

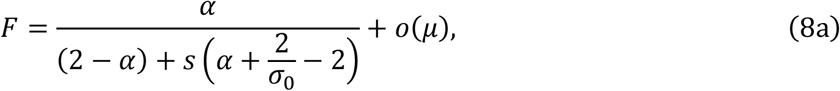

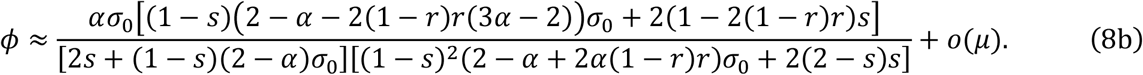

Equations (8a) and (8b) account for the coexistence of outcrossing, selfing and asexual reproduction in a population, and the derivation is available in Section 1.3 of the Mathematica Notebook. *F* converges to the classical expression *F* = *α*/(2 – *α*) as *s* → 0. Note that a high asexual reproduction rate (i.e., large *s*/*σ*_0_) can greatly reduce *F* and *ϕ*, and makes *F* less than 1 even under complete selfing *α* = 1.

Ignoring interactions between deleterious alleles at more than two loci, the genome-wide strength of selection caused by pairwise interactions of deleterious mutations is

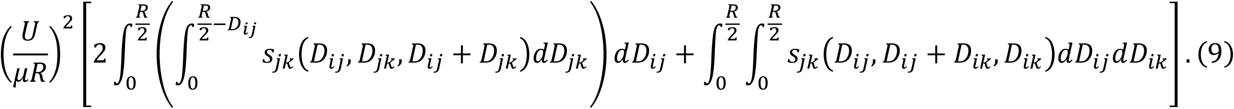

In equation (9), the term *U*/*μR* is the density of fitness loci on the chromosome. The first term in the square bracket corresponds to the contribution from fitness loci located on the same side of the modifier locus, while the second term represents the contribution from fitness loci located on the opposite side of the modifier locus on the chromosome.

#### Hill-Robertson effect

To estimate the strength of selection due to the Hill-Robertson effect, I modify equation (1) in Roze (2021). In a population with a background sexual reproduction rate *σ*_0_ and inbreeding coefficient *F*, the effective recombination rate is *Rσ*_0_(1 – *F*). A modifier that alters the rate of sex by Δ*σ* changes the effective recombination rate by *R*(1 – *F*)Δ*σ*. Given that the effective recombination rate is large compared to selection (i.e., *Rσ*_0_(1 – *F*) ≫ (*h* + (1 – *h*)*F*)*s*), which will not hold when *σ*_0_ → 0 or *F* → 1), the strength of selection caused by the Hill-Robertson effect is

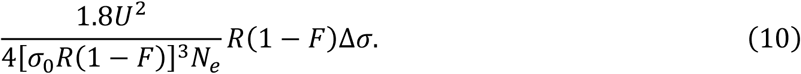

The effective population size *N*_*e*_ is approximately (Glémin 2007)

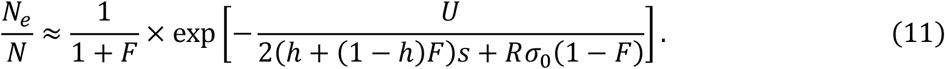

In equation (11), the first term captures the reduction of *N*_*e*_ due to increased variance in allele frequency change under inbreeding (Caballero and Hill 1992). The second term accounts for an increased strength of background selection due to a reduced effective recombination rate under selfing (Nordborg et al. 1996).

### Individual-based simulations

I use individual-based simulations to investigate how selfing influences the overall selection on a sex modifier caused by deleterious mutations in the genome. The simulation is similar to that described in Roze (2015). Briefly, the simulation considers a population with *N* diploid individuals, each carrying two chromosomes with the physical length scaled to be 1. I assume deleterious mutations occur at an infinite number of loci (Kondrashov 1985), and the sex modifier locus is located at the midpoint of the chromosome. At each generation, the number of new deleterious mutations per chromosome is drawn from a Poisson distribution with parameter *U*/2, with their positions on the chromosome being drawn from a uniform distribution *∼* U(0,1). The fitness of an individual is 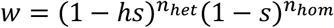, where *n*_*het*_ and *n*_*hom*_ are the number of fitness loci in heterozygous and homozygous state that the individual carries.

To generate the offspring population, for each offspring, I first sample a parent *i* from the parental population, with the probability being proportional to individual fitness *w*_*i*_. I then obtain its rate of sexual reproduction *σ*_*i*_, and generate a random number *ϵ*_1_ *∼* U(0,1). If *ϵ*_1_ > *σ*_*i*_, the offspring is asexually reproduced, and the two chromosomes of the offspring are the same as its parent’s. If *ϵ*_1_ ≤ *σ*_*i*_, the offspring is sexually reproduced, and I draw a second number *ϵ*_2_ *∼* U(0,1) to determine whether the offspring is reproduced from outcrossing or selfing. If *ϵ*_2_ < *α*, the offspring is selfed. If *ϵ*_2_ > *α*, the offspring is outcrossed, and a second parent *j* is sampled with the probability being proportional to *w*_*j*_ (1 – *c*(1 – *σ*_*j*_)), where the term in the parenthesis captures the assumption that asexual reproduction may reduce male gamete production. To obtain gametes generated by meiosis, the number of crossovers between two chromosomes is drawn from a Poisson distribution with parameter *L*, and the position of each crossover is drawn from U(0,1).

For each parameter combination, I simulate the population for 6*N* generations to reach selection-mutation-drift balance, with the sex modifier locus fixed with allele *M*. I then introduce a mutant allele *m* at the sex modifier locus in a randomly chosen chromosome, and run the simulation until *m* becomes fixed or lost. The fixation probability of *m* is calculated based on 2 × 10^5^ replications.

## Results

The result section contains five subsections. The first subsection investigates the effects of selfing on indirect selection caused by intra-locus interactions, assuming that there is no cost of meiosis (set *c* = 1). The second and third subsections present the strength of indirect selection on a sex modifier generated by genome-wide deleterious mutations due to intra-locus interactions and inter-locus interactions, respectively. The fourth subsection presents results from individual-based simulations on how selfing affects the overall selection on a sex modifier caused by deleterious mutations in the genome. The last subsection investigates the effects of selfing on selection due to the cost of meiosis and mate limitation. In this paper, I mainly present results under sporophytic selfing, since results under gametophytic selfing are qualitatively similar. Also, I only present results for a modifier that increases the rate of sex, since when a modifier that increases the rate of sex is favored, a modifier that reduces the rate sex is selected against, and *vice versa*.

### Indirect selection through segregation caused by intra-locus interactions

For indirect selection through segregation, I consider two cases. In the first case, the deleterious allele *a* at the fitness locus is at mutation-selection balance, and I investigate whether a sex modifier allele *m* can invade into a population in which the modifier locus is fixed with allele *M*. In the second case, the fitter allele *A* sweeps from a low initial frequency *p*_*A*,0_ to a high frequency *p*_*A,T*_, and I investigate changes in the frequency of an initially rare sex modifier allele *m*.

#### Mutation-selection balance

Under complete outcrossing or random mating (*α* = 0), a modifier that increases the rate of sex invades when the dominance coefficient of the deleterious mutation *h* is intermediate (Fig. 1(a)). Specifically, when *h* is low 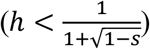, selection causes an excess of *Aa* when there is asexual reproduction (Otto 2003). A modifier that increases the rate of sex enjoys a long-term benefit by breaking this negative association. However, the modifier initially suffers a short-term cost, since the offspring contain more homozygotes, and thus is on average less fit than asexually reproduced offspring. As a result, invasion of the modifier requires that the dominance *h* is not too high, so that the long-term benefit exists, but also not too low so that the short-term cost is not too strong.

**Figure 1.**
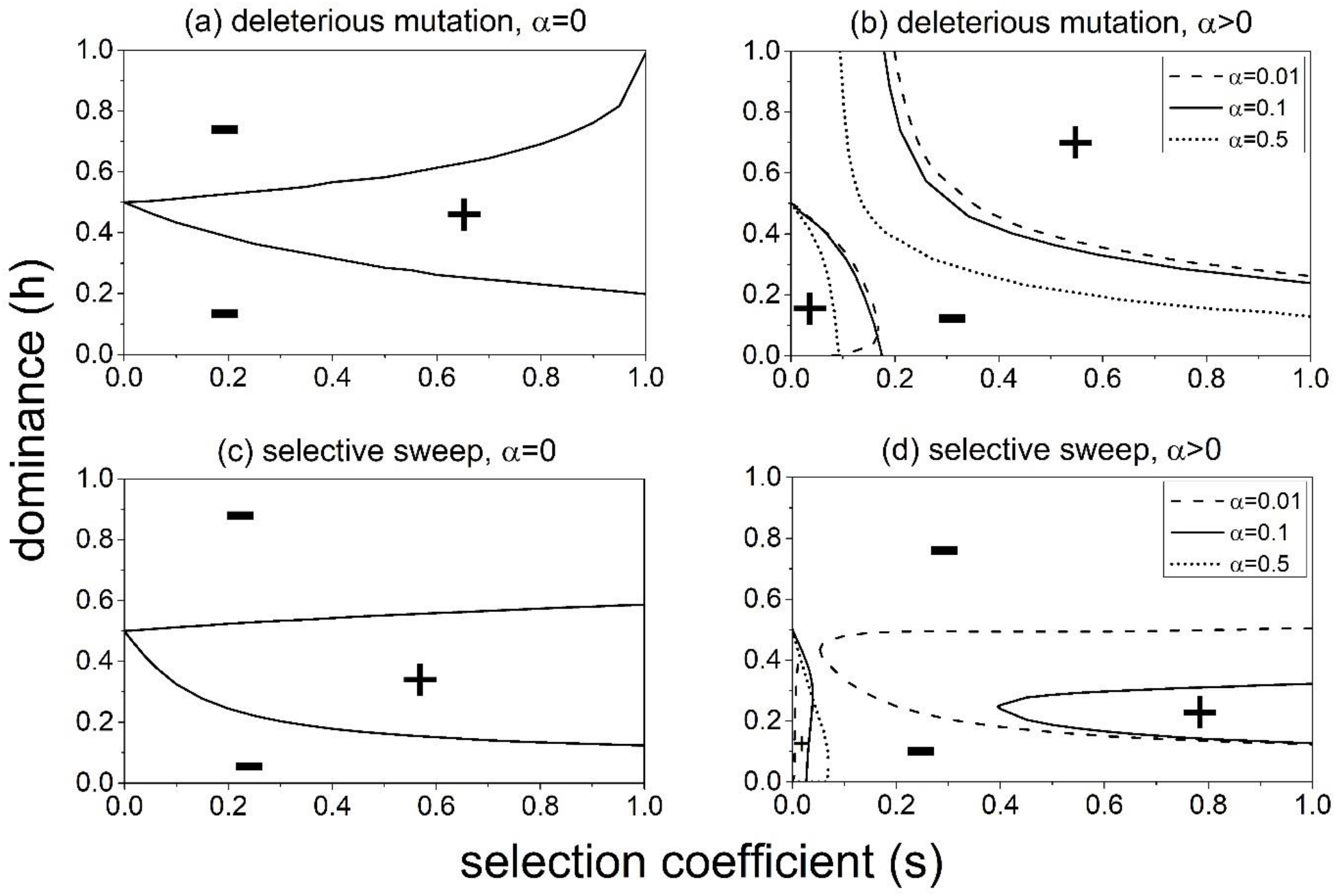
Invasibility of a modifier that increases sexual reproduction when allele *a* at the fitness locus is under selection-mutation balance (panels (a), (b)), and when the fitness locus experiences selective sweep (panels (c), (d)). The symbols “+” and “-” mean that the modifier can invade (or increase in frequency) and cannot invade, respectively. Panels (a) and (c) show results under outcrossing (*α* = 0). Panels (b) and (d) show results when there is selfing. In panel (b), the modifier can invade either at the bottom left or upper right region of the parameter space. In panel (d), the modifier increases in frequency in the top left region or the region between the two lines. For panels (c) and (d), the initial and final frequency of allele *A* is *p*_*A*,0_ = 0.001, *p*_*A,T*_ = 0.999, and the initial frequency of the modifier allele *m* is *p*_*m*,0_ = 0.001. *σ*_0_ = 0.5 for panels (a) and (b), *σ*_0_ = 0.2 for panels (c) and (d). Other parameters are Δ*σ* = 0.01, *h*_*σ*_ = 0.5, *r* = 0.5, *μ* = 10^−5^, *c* = 1, *τ* = 0.The magnitude of the modifier effect Δ*σ* only slightly affects the invasion conditions (Fig. S1).

When selfing is present, given that the selfing rate is much larger than the mutation rate (*α* ≫ *μ*), inbreeding generates an excess of homozygotes at a locus (see equation (8a)). Consequently, the invasion conditions of a modifier increasing the rate of sexual reproduction are altered compared to the case of outcrossing. There exists a narrow range of selection coefficient *s* where the modifier cannot invade for all levels of dominance *h*. This range can be represented by a critical value *θ* under the first-order approximation of the mutation rate *μ* (the expression of *θ* is in Section 3.3 of the Mathematica Notebook). A modifier that increases the rate of sex can invade either when *s* < *θ* and dominance *h* is low enough, or *s* > *θ* and *h* is high enough (bottom-left and upper-right regions in Fig. 1(b)).

The invasion region is mainly determined by the value of the critical selection coefficient *θ*. As *θ* increases, the bottom-left region expands, while the top-right region shrinks. Generally, *θ* becomes smaller when the selfing rate *α* is higher (Fig. 1(b)), the background sexual reproduction rate *σ*_0_ is lower (Figs. 2(a), 2(b)), and when the sex modifier allele *m* is more dominant (compare Figs. 2(a)-(c)). These results are qualitatively similar to those found under gametophytic selfing in Otto (2003). The only difference is that under sporophytic selfing, the critical selection coefficient *θ* will slightly increase when the recombination rate between the modifier and fitness locus *r* is lower (Fig. S2(a)). In contrast, under gametophytic selfing, the value of *θ* is independent of *r* (see equation (13) in Otto (2003)).

**Figure 2.**
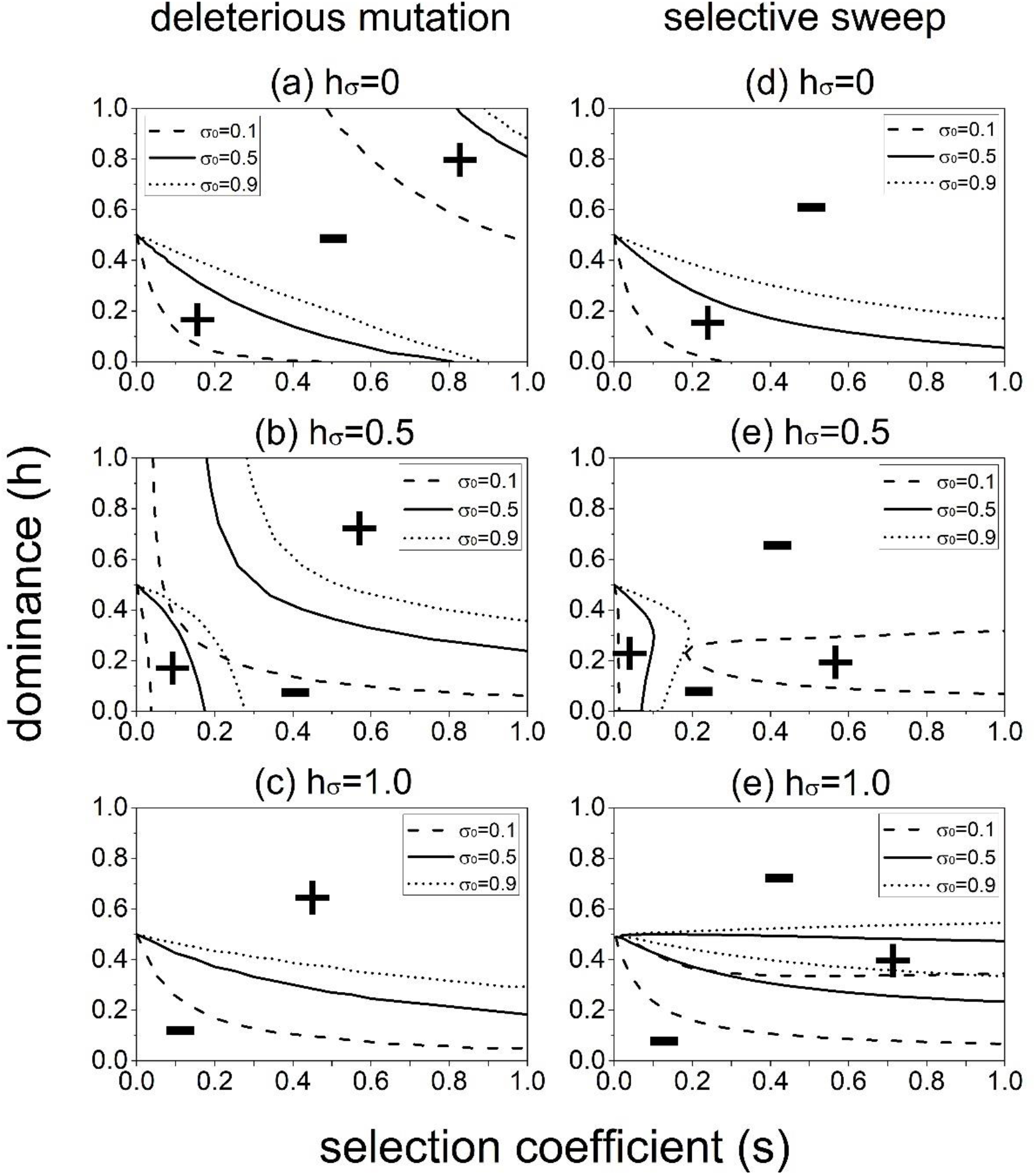
Effects of the dominance coefficient of the modifier (*h*_*σ*_) and background sexual reproduction rate (*σ*_0_) on the invasibility of a modifier that increases sexual reproduction under sporophytic selfing. Results are based on numerical solution of the eigenvalue. Left and right columns show the case when the fitness locus contains deleterious mutations at selection-mutation balance (panels (a)-(c)) and when the fitness locus experiences selective sweep (panels (d)-(f)), respectively. The meanings of “+” and “-” are the same as in Figure 1. For panels (d)-(e), the initial and final frequency of allele *A* is *p*_*A*,0_ = 0.001, *p*_*A,T*_ = 0.999, and the initial frequency of *m* is *p*_*m*,0_ = 0.001. Other parameters are Δ*σ* = 0.01, *α* = 0.1, *r* = 0.5, *μ* = 10^−5^, *c* = 1, *τ* = 0.

We can check the short-term selective (dis)advantage of a sex modifier due to segregation by computing the fitness of offspring from sexual and asexual reproduction. Section 1.4 of the Mathematica Notebook Materials shows that the relative fitness reduction of asexually reproduced and selfed offspring compared to outcrossed offspring is respectively,

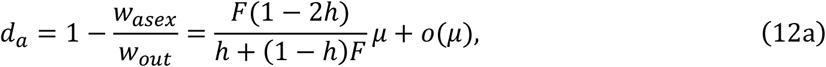

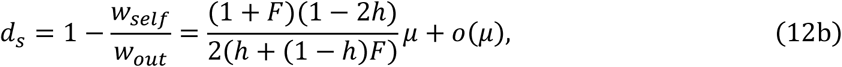

where *F* is given by equation (8a), and *d*_*s*_ is just inbreeding depression. Equations (12a) and (12b) imply that when the deleterious allele is partially recessive (h < 0.5), both asexually reproduced and selfed offspring are less fit than outcrossed offspring. Since 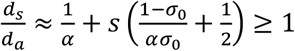, selfed offspring are less fit than asexually reproduced offspring. Moreover, Section I in Supplementary Materials shows that a lower background sexual reproduction rate *σ*_0_ or selfing rate *α* will increase the relative fitness of asexually reproduced offspring *d*_*a*_, but reduce the relative fitness of selfed offspring *d*_*s*_ (i.e., stronger inbreeding depression).

Since sexually reproduced offspring contain both selfed and outcrossed offspring, their mean fitness is *w*_*sex*_ = *α*(1 – *d*_*s*_) + 1 – *α* = 1 – *αd*_*s*_. When the fitness locus is under selection-mutation balance the fitness reduction of sexually versus asexually reproduced offspring is

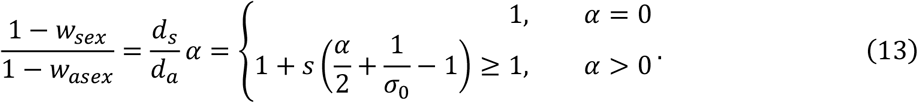

When deleterious alleles are partially recessive (h < 0.5), as *w*_*sex*_, *w*_*asex*_ < 1, equation (13) shows that sexually reproduced offspring are less fit than asexually reproduced offspring. This disadvantage of sexual reproduction is greater under a higher selfing rate (*α*) and a lower background sexual reproduction rate (*σ*_0_), indicating a stronger short-term fitness cost of modifiers that enhance sexual reproduction. When *h* > 0.5, as *w*_*sex*_, *w*_*asex*_ > 1, sexually reproduced offspring are fitter than asexually reproduced offspring, and this advantage increases with higher *α* and lower *σ*_0_.

The above results qualitatively hold under gametophytic selfing. However, unlike sporophytic selfing, the fitness of sexually reproduced offspring relative to asexually reproduced offspring is independent of the gametophytic selfing rate (see section II in Supplementary Materials).

#### Selective Sweep

When allele *A* sweeps from a low to high frequency at the fitness locus, selfing generally inhibits the evolution of sexual reproduction. Under outcrossing, similar to the results under mutation-selection balance, a modifier increasing the sexual reproduction rate increases in frequency when the dominance *h* is intermediate (Fig. 1(c)). When selfing is present, this region continuously shrinks as the selfing rate *α* increases (region Fig. 1(d)). However, there appears a new parameter region that the modifier will increase in frequency, which occurs when *s* is small and dominance *h* is low (bottom-left corner in Fig. 1(d)). This region expands when the selfing rate becomes higher (Fig. 1(d)) and the sex modifier *m* is more recessive (compare Figs. 2(d)-(e)). Moreover, a higher recombination rate *r* slightly reduces the parameter regions that favor a higher sexual reproduction rate (Fig. S2(b)).

#### Genetic association analysis

Selfing generates positive association between homozygotes at different loci (see equation (8a)). Therefore, genotype *mm* will be associated with homozygotes at the fitness locus (*AA* and *aa*), while *Mm* will be associated with heterozygote *Aa*. When *h* is low so that *Aa* is on average fitter than homozygotes 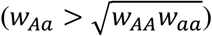, the modifier *m* found in *Mm* individuals will enjoy a short-term benefit by associating with *Aa*, but suffers long-term disadvantages due to a reduction in genetic variance. In contrast, *m* found in *mm* individuals is associated more with homozygotes *AA* and *aa*, thereby suffering a short-term cost but enjoying a long-term benefit. The invasion of *m* found in *Mm* individuals requires a low enough *h* so that the short-term benefit is strong enough, while the invasion of *m* found in *mm* individuals requires a high enough *h*.

Under mutation-selection balance, consistent with Otto (2003), when selection is weak (*s* < *θ*), the evolution of the modifier is mainly determined by those found in *Mm* individuals, so allele *m* is associated more with *Aa* (i.e., *corr*(*m, Aa*) > 0). Therefore, invasion occurs when *h* is low enough. When selection is strong (*s* > *θ*), the evolution is mediated by *m* often found in *mm* individuals (thus *corr*(*m, Aa*) < 0), so invasion occurs when *h* is high enough.

Under selective sweep, when selection is weak, the modifier *m* is associated with *Aa*, so a sex modifier *m* increases in frequency when *h* is low enough. When selection is strong, whether the correlation between the modifier *m* and the heterozygote *Aa* will be positive or negative depends on the dominance *h*. When *h* is low, the modifier *m* is associated with homozygotes, so the modifier *m* is favored when *h* is high enough. As *h* increases, the modifier *m* becomes associated with *Aa*, so the modifier *m* is favored when *h* is low enough. As a result, the modifier will increase in frequency only when *h* is intermediate, and this intermediate range may not exist for some values of selection coefficient.

### Strength of indirect selection through segregation caused by intra-locus interactions

The previous section investigates selection contributed by a single fitness locus. This result can be extended to estimate the strength of indirect selection on a modifier of sex through segregation generated by deleterious mutations in the genome. When a modifier allele is rare and has small effects, the selective strength Δ*p*_*m*_/*p*_*m*_(1 – *p*_*m*_) can be approximated by the leading eigenvalue as *λ* – 1 (Otto (2003)). For simplicity, I assume loci are identical and ignore genetic associations between loci and epistasis. When the genomic mutation rate is *U*, the number of identical fitness loci is *L* = *U*/*μ*, and the genome-wide strength of selection is approximately *L*(*λ* – 1). In addition, I estimate the strength of the short-term (dis)advantage of a sex modifier due to fitness differences between sexually and asexually reproduced offspring caused by deleterious mutations in the genome. When there are *L* loci, the fitness of asexually reproduced and selfed offspring relative to outcrossed offspring is 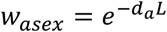 and 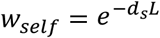, where *d*_*a*_ and *d*_*s*_ are given by equation (12). The strength of this short-term selective force can thus been estimated by setting 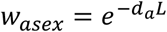 and 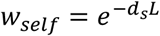 in equation (6), and solving for the leading eigenvalue.

Previous analyses show that there are two major categories of deleterious mutations: small-effect, partially dominant mutations and large-effect, highly recessive lethal mutations (Mukai et al. 1972, Eyre-Walker and Keightley 2007, Charlesworth and Willis 2009). Based on previous estimations (Simmons and Crow 1979), I compare the strength of indirect and direct selection caused by deleterious mutations by setting *s* = 0.05, *h* = 0.2 for small-effect deleterious mutations, and *s* = 0.8, *h* = 0.02 for large-effect lethal mutations.

For selection caused by small-effect deleterious mutations, a modifier that increases the rate of sex is indirectly selected for either when the rate of sexual reproduction *σ*_0_ is not low and the selfing rate is not high (region I in Fig. 3(a)), or when *σ*_0_ is low and *α* is high (region II in Fig. 3(a)). In region I, sexually reproduced offspring is fitter than asexually reproduced offspring, so the modifier enjoys a weak short-term advantage (upper-left corner in Fig. 3(b)). In contrast, in region II, fitness of sexually reproduced offspring is much lower than asexually reproduced offspring, indicating a strong short-term cost to the modifier (bottom-right corner in Fig. 3(b)). However, the modifier is actually strongly favored by indirect selection (region II in Fig. 3(a)), which suggests that the modifier has a strong long-term benefit by increasing the equilibrium mean fitness in the population.

**Figure 3.**
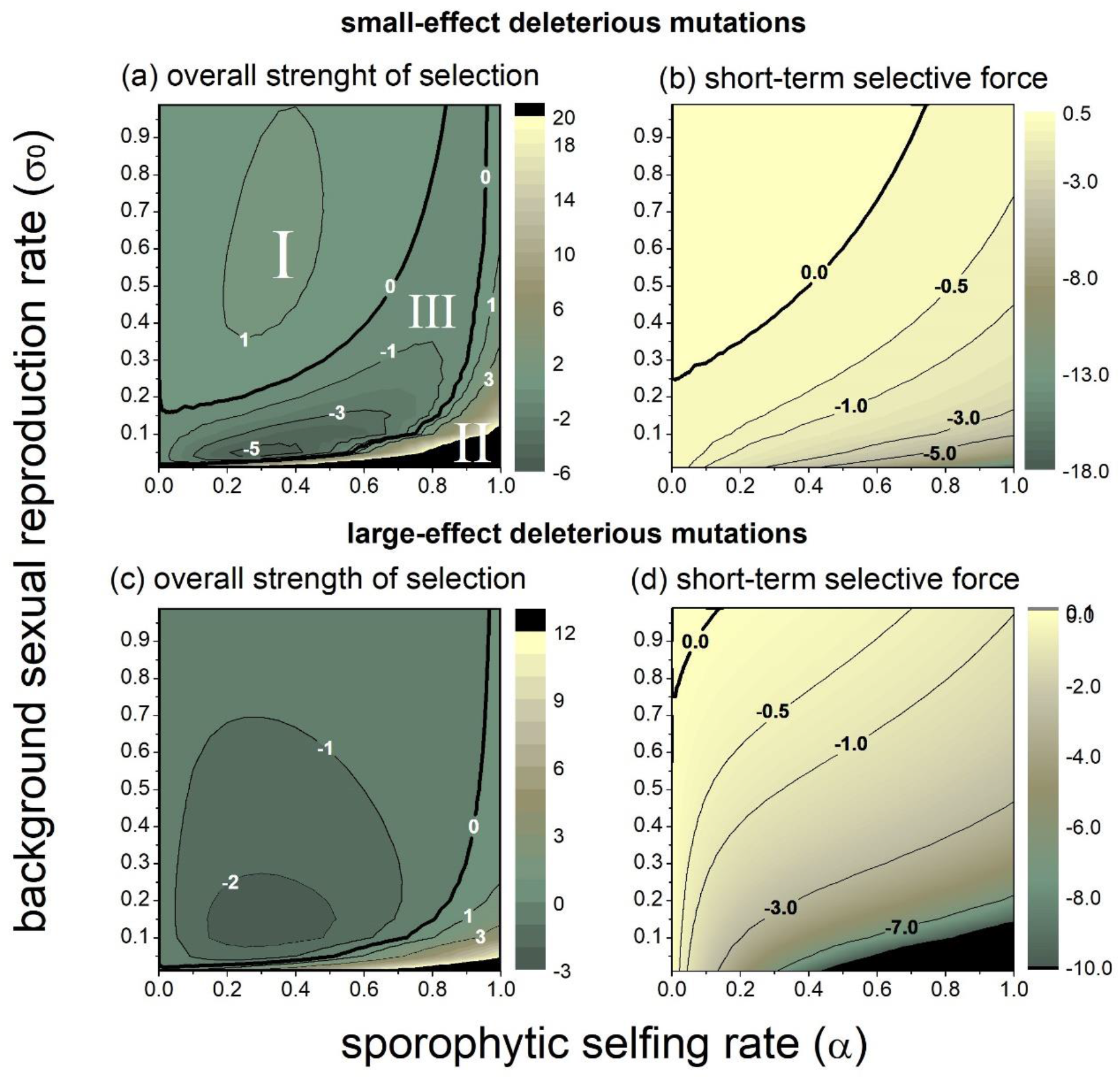
Genome-wide strength of indirect selection (in the order of 10^−4^) on a modifier increasing the sexual reproduction rate caused by within-locus interactions. The upper and bottom panels show selection caused by small-effect and large-effect deleterious mutations. Panels (a) and (c) show the overall strength of indirect selection. Panels (b) and (d) show the strength of short-term (dis)advantage of the modifier due to fitness differences between sexually and asexually reproduced offspring. Similar results are found under gametophytic selfing (Fig. S3). I use *s* = 0.05, *h* = 0.2, *U* = *Lμ* = 0.2 for small-effect deleterious mutations, and *s* = 0.8, *h* = 0.02, *U* = *Lμ* = 0.025 for large-effect deleterious mutations (Mukai et al. 1972, Simmons and Crow 1977, Klekowski and Godfrey 1989). Other parameters are Δ*σ* = 0.01, *h*_*σ*_ = 0.5, *c* = 1, *τ* = 0, *r* = 0.5.

I also investigated the influences of the selection coefficient and dominance of deleterious mutations. Consistent with the results in the previous subsection, a modifier increasing the rate of sexual reproduction is more likely to invade when the selection coefficient of small-effect deleterious mutations is smaller (Figs. S4(a)-(b)). Also, a lower *h* reduces the relative fitness of sexually reproduced offspring relative to asexually reproduced offspring, and thus reduces the parameter space for the modifier to invade (Figs. S4(c)-(d)).

For selection generated by large-effect, highly recessive mutations, a modifier increasing the rate of sexual reproduction is indirectly selected against in a large parameter space (Fig. 3(c)). This is partly contributed by the short-term disadvantage of the modifier, as sexually reproduced offspring tend to have lower fitness than asexually reproduced offspring when deleterious alleles are very recessive (see Fig. 3(d)). However, similar to selection generated by small-effect deleterious mutations in Fig. 3(a), the modifier is favored when the background sexual reproduction rate *σ*_0_ is low and the selfing rate *α* is high (bottom-right corner in Fig. 3(c)).

Regarding selection on a sex modifier when both categories of deleterious mutations are present in the genome (compare Figs. 3(a) and 3(c)), the only parameter space where the modifier will always be favored is when the background sexual reproduction rate *σ*_0_ is low, and the selfing rate *α* is high (bottom-left corner in panels of Fig. 3(a), 3(c)). On the other hand, in the parameter space represented by region III in Fig. 3(a), the modifier should always be selected against. When *σ*_0_ is relatively high, and *α* is relatively low (region I in Fig. 3(a)), small-effect deleterious mutations and highly recessive, large-effect deleterious mutations have opposing effects on the evolution of sex with comparable selective strength. Therefore, in this parameter region, whether a modifier that increases the rate of sexual reproduction is favored or not should depend on the relative genomic mutation rate of the two categories of mutations.

### Strength of indirect selection caused by inter-locus interactions

Here I estimate how selfing impacts the strength of indirect selection generated by deleterious mutations in the genome due to inter-locus interactions. This indirect selection can be caused by two forces: deterministic interactions between deleterious mutations, and the Hill-Robertson effect in a finite population (Stetkenko and Roze 2022). Generally, unless the selfing rate is high, indirect selection on a sex modifier caused by inter-locus interactions is weak compared to indirect selection caused by intra-locus interactions, partly consistent with findings in Roze and Michod (2010) for outcrossing populations.

#### Deterministic interactions between deleterious mutations

As Fig. 4(a) illustrates, even with a small map length (*R* = 1), indirect selection caused by deterministic interactions between deleterious mutations is quite weak. Selfing causes selection against sex when there is no epistasis (solid lines in Fig. 4(a)), but can favor sex when there is negative epistasis between deleterious mutations (dashed lines in Fig. 4(a)). Generally, the background sexual reproduction rate *σ*_0_ only slightly influences how the strength of selection changes with the selfing rate, except when the selfing rate is high. Under complete selfing, the strength of selection will reduce to 0 in a completely sexually reproducing population since the inbreeding coefficient becomes *F* = 1 (Stetsenko and Roze 2022). In contrast, when there is asexual reproduction, the strength of selection will not reduce to 0 under complete selfing (see results at *α* = 1 in Fig. 4(a)), because *F* remains smaller than 1 (see equation (8a)).

**Figure 4.**
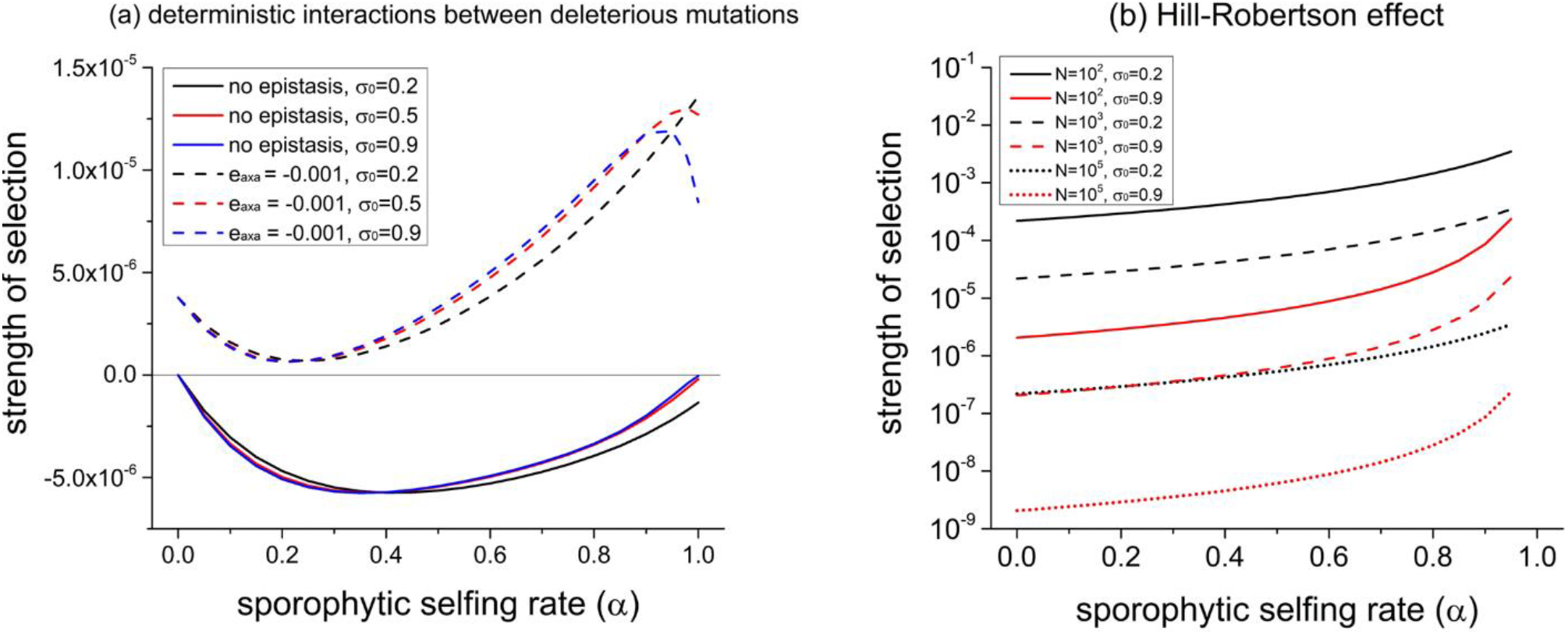
Strength of indirect selection (Δ*p*_*m*_/*p*_*m*_(1 – *p*_*m*_)) on a rare sex modifier due to effective changes in the recombination rate. Panels (a) and (b) show the strength of selection caused by deterministic interactions between deleterious mutations, and the Hill-Robertson effect in a finite population, respectively. In panel (a), solid lines show results when fitness loci have multiplicative interactions on individual fitness (no epistasis). Dashed lines show results under additive-additive epistasis, so that for two loci *j* and *k*, the fitness of genotype *A*_*j*_*a*_*j*_*A*_*k*_*a*_*k*_, *A*_*j*_*a*_*j*_*a*_*k*_*a*_*k*_, and *a*_*j*_*a*_*j*_*a*_*k*_*a*_*k*_ is further changed from the multiplication by *e*_a×a_, 2*e*_a×a_, and 4*e*_a×a_, respectively. Other parameters are *s* = 0.05, *h* = 0.2, *U* = 0.2, Δ*σ* = 0.01, *h*_*σ*_ = 0.5, *R* = 1.

#### The Hill-Robertson effect

Fig. 4(b) shows that selection on a sex modifier due to the Hill-Robertson is weak compared to selection caused by intra-locus interactions (*∼* 10^−4^ – 10^−3^), unless the population size is small (*N ∼* 10^2^), the background sexual reproduction rate *σ*_0_ is low and the selfing rate *α* is high (black solid line in Fig. 4(b)). As equations (13) and (14) indicate, selfing increases the strength of selection due to the Hill-Robertson effect by reducing both the effective recombination rate and effective population size, while a higher background sexual reproduction rate *σ*_0_ reduces the selective strength, as illustrated in Fig. 4(b).

### Overall effects of selfing on indirect selection caused by deleterious mutations

Results from Individual-based simulations in Fig. 5 confirm that selection on a sex modifier is mainly generated by intra-locus interactions rather than inter-locus interactions, unless the selfing rate is high (but see Fig. 4 in Roze and Michod (2010) for results when deleterious alleles are weakly recessive under outcrossing). Considering selection generated by small-effect deleterious mutations, a modifier that enhances sexual reproduction is generally disfavored when the background sexual reproduction rate *σ*_0_ is low, unless the selfing rate is high (line with *σ*_0_ = 0.2 in Fig. 5(a)). Under a higher level of *σ*_0_, as the selfing rate increases, the fixation probability of the sex modifier is reduced within a specific range of the selfing rate (see the range of *α*=0.6-0.9 at *σ*_0_ = 0.5, 0.9 in Fig. 5(a)). For selection caused by highly recessive, large-effect alleles, the modifier tends to be strongly disfavored under an intermediate selfing rate, unless the selfing rate is high (Fig. 5(b)). Referring to the results at *σ*_0_ = 0.2, 0.5, 0.9 in Fig. 3(a) and 3(c), the above findings are qualitatively align with results for indirect selection through segregation caused by intra-locus interactions.

**Figure 5.**
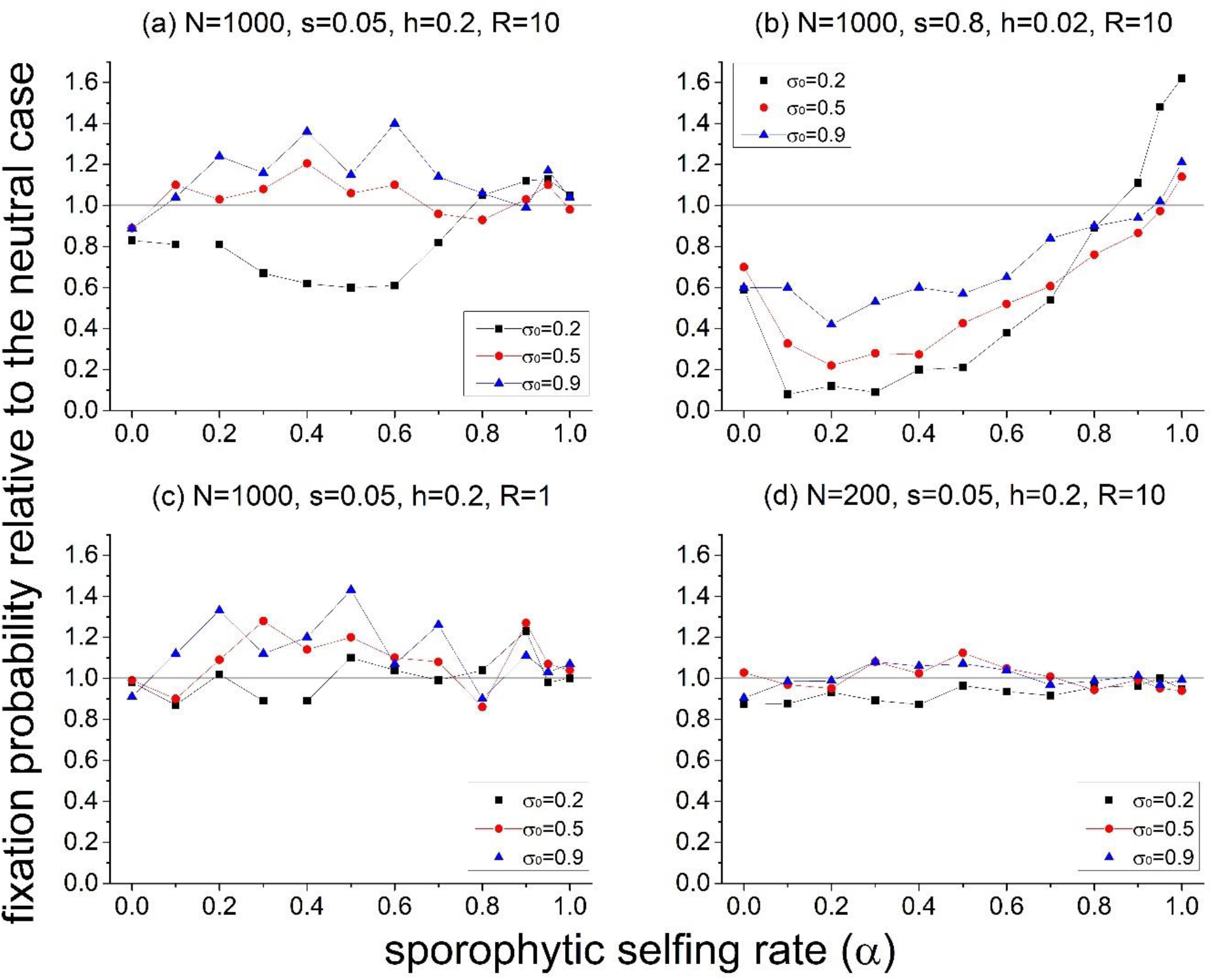
Effects of selfing on the fixation probability of a sex modifier allele with deleterious mutations from individual-based simulations. The fixation probability is divided by that under the neutral case 1/2*N*. The haploid genomic mutation rate is *U* = 0.2 for partially recessive, small-effect deleterious mutations (panels (a), (c), (d)), and *U* = 0.05 for highly recessive, large-effect mutations (panel (b)). Other parameters are Δ*σ* = 0.05, *h*_*σ*_ = 0.5, *c* = 1, *τ* = 0.

Moreover, when the genomic recombination rate is low or the population size is small, selection driven by intra-locus interactions still tends to be the dominant force, despite the increased strength of selection caused by inter-locus interactions (Figs. 5(c), 5(d)). When the background sexual reproduction rate *σ*_0_ is low, a small map length or a small population size weakens selection against the sex modifier (compare results at *σ*_0_ = 0.2 in Figs. 5(c), 5(d) with 5(a)). Nevertheless, this effect may primarily stem from a reduced short-term cost of sex caused by intra-locus interactions. Specifically, under tight linkage or a small population size, there will be fewer segregating deleterious mutations, which can increase the fitness of sexually reproduced offspring relative to asexually reproduced offspring (Kirkpatrick and Batillon 2000). Moreover, under a higher level of *σ*_0_ (*σ*_0_ = 0.5,0.9), the pattern of how the fixation probability changes with the selfing rate *α* in Fig. 5(c) and 5(d) similar to Fig. 5(a).

Moreover, a high selfing rate can increase the strength of selection on a sex modifier by lowering the effective recombination rate, it also reduces the effective population size (see equation (10)). As a result, under a high selfing rate, the fixation probability is close to the neutral case when deleterious mutations are partially dominant and have small effects (Figs. 5(a), (c), (d)).

### Selection caused by the cost of meiosis and mate limitation

Previous subsections focus on indirect selection on a sex modifier caused by selection at fitness loci. To quantify how selfing can reduce the strength of selection against sex by lowering the cost of meiosis and providing reproductive assurance under mate limitation, I define the strength of mate limitation *τ* as the reduced proportion of outcrossed ovules (Xu 2022). Therefore, the relative fertility via outcrossing is 1 – *τ* compared to selfing and asexual reproduction. The strength of selection on a sex modifier can be obtained by setting *s* = 0 and *w*_*asex*_ = *w*_*self*_ = 1/(1 – *τ*) in the recursion equation (6). For a rare modifier allele with small effect (|Δ*σ*| ≪ 1), the selective strength is approximately *s*_*e*_(*h*_*σ*_ + (1 – *h*_*σ*_)*F*), where the effective selection coefficient *s*_*e*_ is

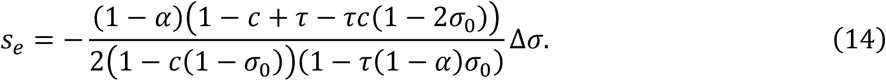

In the denominator, the term 2(1 – *c*(1 – *σ*_0_)) is the amount of male gamete production, and the term 1 – *τ*(1 – *α*)*σ*_0_ accounts for fitness reduction due to mate limitation. In the numerator, the term 1 – *c* and *τ* capture selection due to the cost of meiosis and mate limitation, respectively, while the –*τc*(1 – 2*σ*_0_) captures interactions between the two forces.

Fig. 6 shows that generally, the strength of selection caused by the cost of meiosis and mate limitation is often comparable to indirect selection generated by deleterious mutations in the genome. Interestingly, the strength of selection against the modifier is stronger when the background sexual reproduction rate *σ*_0_ is higher (compare solid and dashed lines in Fig. 6). This is because when there is mate limitation, a higher *σ*_0_ reduces the mean fitness of the population (the term 1 – *τ*(1 – *α*)*σ*_0_ in equation (14)). Generally, a higher selfing rate tends to reduce the strength of selection. However, under strong mate limitation (*τ* = 0.8 in Fig. 6), selection against sexual reproduction is strongest under a low selfing rate rather than under complete outcrossing. This is because although selfing reduces the effective selection coefficient *s*_*e*_, it strengthens selection by increasing homozygosity (the term *h*_*σ*_ + (1 – *h*_*σ*_)*F*).

**Figure 6.**
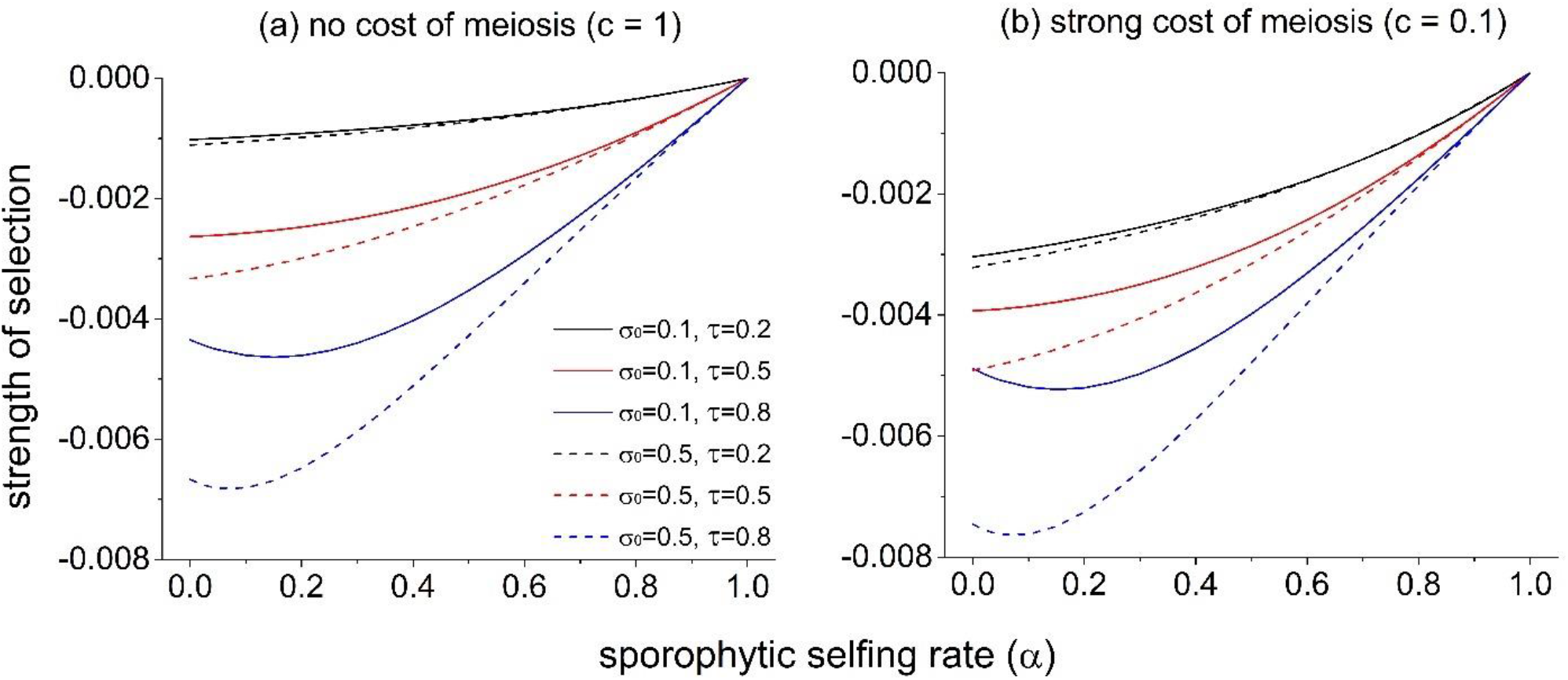
Effects of the selfing rate on the strength of selection against a modifier that increases sex (Δ*σ* = 0.01, *h*_*σ*_ = 0.5) caused by the cost of meiosis and mate limitation. Panels (a) and (b) show results when there is no and strong cost of meiosis, respectively. Lines with different colors show results under different strength of mate limitation. The strength of selection is calculated based on equation (14).

## Discussion

This study investigates the effects of selfing on the evolution of an allele through three major selective forces: (1) indirect selection caused by an altered rate of segregation at selected loci, referred to as selection caused by intra-locus interactions, (2) indirect selection due to changes in the effective recombination rate between selected loci, referred to as selection caused by inter-locus interactions, and (3) selection due to the cost of meiosis and mate limitation. The effects of selfing on the three selective forces are summarized in Table 1. Briefly, selfing affects indirect selection caused by both intra-locus and inter-locus interactions by primarily increasing homozygosity across the genome, and generating correlation between homozygotes at different loci, as indicated by equations (8a) and (8b). Also, similar to asexual reproduction, selfing is also a mode of uniparental reproduction. Therefore, selfing can weaken, selection against sexual reproduction due to the cost of meiosis and mate limitation, although may not revert the direction.

The mechanisms by which selfing influences indirect selection caused by intra-locus selection are complicated. One important mechanism is that selfing will induce higher homozygosity in sexually reproduced offspring compared to asexually reproduced offspring. Therefore, when deleterious alleles are recessive, similar to inbreeding depression, sexually reproduced offspring will be less fitness than asexually reproduced offspring, resulting in a short-term disadvantage of a modifier that increases the rate of sexual reproduction. Selfing also affects indirect selection by altering the genetic associations between a modifier locus and fitness loci. Generally, under sporophytic selfing, a modifier that increases the rate of sexual reproduction can invade in two scenarios: either when the deleterious allele is recessive enough under weak selection, or when the allele is dominant enough under stronger selection. These results are qualitatively similar to the findings under gametophytic selfing (Otto 2003).

Consequently, unless the selfing rate is high, highly recessive, large-effect deleterious mutations in the genome often exert indirect selection against a modifier that enhances sexual reproduction through intra-locus interactions. In contrast, partially recessive, small-effect deleterious mutations can favor the evolution of sexual reproduction in a larger parameter space. In general, a modifier that enhances sexual reproduction will be indirectly selected against by intra-locus interactions at an intermediate selfing rate and when the sexual reproduction rate in the population is low. Given that the map length in most organisms lies between 1 ∼ 10 (Otto and Payseur 2019), indirect selection caused by inter-locus interactions tends to be much weaker than indirect selection caused by intra-locus interactions, unless deleterious mutations are weakly recessive and the population is random mating (Roze and Michod 2010). However, when the selfing rate is very high, both intra-locus and inter-locus interactions will generate strong indirect selection for a modifier that increases the rate of sexual reproduction.

By comparing the strength of the three selective forces on a sex modifier, I find that the dominant selective force will depend on the rates of selfing and sexual reproduction. Under outcrossing, selection on a sex modifier is primarily driven by the cost of meiosis and mate limitation. Deleterious mutations in the genome result in weak indirect selection against a higher sexual reproduction rate. As the selfing rate increases, selection due to the cost of meiosis and mate limitation gets weaker, while indirect selection through intra-locus interactions becomes stronger and may become the dominant force. In highly selfing populations, indirect selection caused by both intra-locus and inter-locus interactions is significant. Furthermore, under a higher rate of sexual reproduction, selection on a sex modifier due to the cost of meiosis and mate limitation intensifies, while indirect selection generated by deleterious mutations weakens. This suggests that evolutionary transition from complete sexual reproduction to partially asexual reproduction may initially be driven by the cost of meiosis and mate limitation.

The current results suggest that high selfing rate should favor the evolution of sexual reproduction is consistent with the evidence that populations with high selfing rates are rarely found in partially clonal herbs (Vallejo-Marín et al. 2010). Empirical evidence show that apomictic populations often derive from self-incompatible populations (Hörandl 2010, Hojsgaard and Hörandl 2019). However, our results suggest that intermediate selfing rates tend to favor the evolution of asexual reproduction, suggesting that asexual reproduction tends to evolve before, instead of after, the evolution of selfing. In fact, equation (12b) shows that asexual reproduction will inhibit the evolution of selfing by increasing inbreeding depression. This hypothesis may be tested based on ancestral state reconstructions.

The offers a novel interpretation for the observed association between vegetative reproducing populations and self-incompatibility/outcrossing (Ellstrand and Roose 1987, Vallejo-Marín et al. 2010). Traditionally, it has been proposed that in partially clonal populations, outcross-pollination among cloned individuals (geitonogamy) will produce offspring equivalent to those produced through selfing, thus suffering the cost of inbreeding depression (Vallejo-Marín et al. 2010). Consequently, mechanisms avoiding geitonogamy may be favored, such as self-incompatibility and dichogamy (temporal separation of female and male function). However, I find that assuming no spatial structure, even with geitonogamy, the fitness of outcrossed offspring is always higher than of asexually reproduced offspring (equation (12a)). Moreover, equation (12b) reveals that a higher level of asexual reproduction can lower the fitness of selfed offspring, potentially selecting against alleles that increases the rate of selfing. Thus, in partially clonal populations, mechanisms promoting obligate outcrossing may be primarily selected for by avoiding intra-individual self-fertilization, instead of geitonogamy.

This study did not explicitly model indirect selection on sex for segregation when departure from Hardy-Weinberg equilibrium is generated by genetic drift (Balloux et al. 2003). Specifically, in a finite population, an oversampling of heterozygotes brings the allele frequency closer to 0.5, and thus will persist longer than an oversampling of homozygotes, which brings the allele frequency closer to fixation. Under random mating, this effect can thus lead to an excess of heterozygosity, which can favor sex when deleterious mutations are weakly recessive or dominant (Roze and Michod 2010). Roze and Michod (2010) also suggest that the strength of selection on sex for segregation generated by drift is comparable to that generated by selection at each locus. However, under inbreeding, selection generated by drift should be much weaker than the deterministic force generated by selection at each locus, since departure from Hardy-Weinberg equilibrium will be dominated by the interaction between inbreeding and selection. In fact, under selfing, the selective strength on sex generated by selection at each locus is 1/*μ* time stronger than under outcrossing, where *μ* is the mutation rate of deleterious alleles (Otto 2003).

Also, the current study did not explore the impact of selfing on the evolution of sex due to the cost of males. However, empirical evidence suggests that selfing may generally weaken the cost of males by reducing resource allocation to the male function in plants. Higher selfing rates are correlated with lower pollen-ovule ratios (Cruden 1977, Cruden and Lyon 1985, de Cunha and Aizen 2023) and smaller flower display size (Sicard et al. 2011). While a corresponding increase in allocation to the female function might be anticipated, the evidence is less conclusive (Cruden 1977, Cruden and Lyon 1985). In some groups, selfing species produce more ovules per flower than outcrossing species (Moine and Anderson 1992, Delesalle et al. 2008). In other groups, selfing reduces the ovule number per flower (Ritland and Ritland 1989). This may be because in highly selfing populations, male and female gamete production is expected to be perfectly correlated (Mazer et al. 2007). Also, selfing populations may experience smaller stochastic variation in mating success, which selects for fewer ovules (Burd et al. 2009). Ovule production may not fully reflect female function allocation. In fact, self-compatible species tend to have a high fruit-flower ratio than self-incompatible species, indicating a greater allocation to the female function (Sutherland 1986).

The current study investigates the evolution of sexual reproduction through selection within a population. Another category of studies on the evolution of sex looks at selection between populations, by investigating how the sexual reproduction rate affects the mean fitness of a population (reviewed in Otto (2009)). Analyses in Section I of Supplementary Materials suggest that selfing tends to increase the strength of between-population selection that favors sexual reproduction.

In conclusion, this study indicates that an intermediate selfing rate tends to impede the evolution of sexual reproduction, especially when the baseline sexual reproduction rate is low. A high selfing rate strongly favors the evolution of sexual reproduction. Regarding empirical evidence, these findings suggest that the evolution of asexual reproduction may often precede the evolution of selfing.

## Acknowledgments

I would like to thank the editor Dr. Tim Connallon, the associate editor Dr. Pierre-Olivier Cheptou, two anonymous reviewers, Dr. Maria Servedio, and Brian Lerch for their help comments on the manuscript. I also thank Dr. Sally Otto for her help with identifying an error in a previous version of the model.

## References

Agrawal, A. F., & Chasnov, J. R. (2001). Recessive mutations and the maintenance of sex in structured populations. Genetics, 158(2), 913–917.

Asker, S. E., & Jerling, L. (2017). Apomixis in plants. Routledge.

Balloux, F., Lehmann, L., & de Meeûs, T. (2003). The population genetics of clonal and partially clonal diploids. Genetics, 164(4), 1635–1644.

Barrett, S. C. (2008). Major evolutionary transitions in flowering plant reproduction: an overview. International Journal of Plant Sciences, 169(1), 1–5.

Barrett, S. C. (2015). Influences of clonality on plant sexual reproduction. Proceedings of the National Academy of Sciences, 112(29), 8859–8866.

Barton N. H. (1995). A general model for the evolution of recombination. Genetic Research, 65, 123–144.

Bell, G. (1982). The paradox of sexuality. The masterpiece of nature: The evolution and genetics of sexuality, 19–78.

Bengtsson, & Ceplitis. (2000). The balance between sexual and asexual reproduction in plants living in variable environments. Journal of Evolutionary Biology, 13(3), 415–422.

Burd, M., Ashman, T. L., Campbell, D. R., Dudash, M. R., Johnston, M. O., Knight, T. M., … & Vamosi, J. C. (2009). Ovule number per flower in a world of unpredictable pollination. American Journal of Botany, 96(6), 1159–1167.

Busch, J. W., & Delph, L. F. (2012). The relative importance of reproductive assurance and automatic selection as hypotheses for the evolution of self-fertilization. Annals of botany, 109(3), 553–562.

Caballero, A., & Hill, W. G. (1992). Effective size of nonrandom mating populations. Genetics, 130(4), 909–916.

Charlesworth, B. (1980). The cost of sex in relation to mating system. Journal of Theoretical Biology, 84(4), 655–671.

Charlesworth, B. (2013). Background selection 20 years on: the Wilhelmine E. Key 2012 invitational lecture. Journal of Heredity, 104(2), 161–171.

Charlesworth, B., Charlesworth, D., & Morgan, M. T. (1990). Genetic loads and estimates of mutation rates in highly inbred plant populations. Nature, 347(6291), 380–382.

Charlesworth, D., & Willis, J. H. (2009). The genetics of inbreeding depression. Nature reviews genetics, 10(11), 783–796.

Charlesworth, D., Charlesworth, B., & Strobeck, C. (1979). Selection for recombination in partially self-fertilizing populations. Genetics, 93(1), 237–244.

Chasnov, J. R. (2000). Mutation-selection balance, dominance and the maintenance of sex. Genetics, 156(3), 1419–1425.

Cruden, R. W. (1977). Pollen-ovule ratios: a conservative indicator of breeding systems in flowering plants. Evolution, 32–46.

Cruden, R. W., & Lyon, D. L. (1985). Patterns of biomass allocation to male and female functions in plants with different mating systems. Oecologia, 66, 299–306.

da Cunha, N. L., & Aizen, M. A. (2023). Pollen production per flower increases with floral display size across animal-pollinated flowering plants. American Journal of Botany, e16180.

Damgaard, C., & Abbott, R. J. (1995). Positive correlations between selfing rate and pollen-ovule ratio within plant populations. Evolution, 214–217.

Damgaard, C., & Loeschcke, V. (1994). Genotypic variation for reproductive characters, and the influence of pollen-ovule ratio on selfing rate in rape seed (Brassica napus). Journal of Evolutionary Biology, 7(5), 599–607.

Delesalle, V. A., Mazer, S. J., & Paz, H. (2008). Temporal variation in the pollen: ovule ratios of Clarkia (Onagraceae) taxa with contrasting mating systems: field populations. Journal of Evolutionary Biology, 21(1), 310–323.

Eckert, C. G. (1999). Clonal plant research: proliferation, integration, but not much evolution. American Journal of Botany. 86(11), 1649–1654.

Eckert, C. G., Samis, K. E., Dart, S., Harder, L. D., & Barrett, S. C. H. (2006). Reproductive assurance and the evolution of uniparental reproduction in flowering plants. Ecology and evolution of flowers, 183, 203.

Ellstrand, N. C., & Roose, M. L. (1987). Patterns of genotypic diversity in clonal plant species. American Journal of Botany, 74(1), 123–131.

Eyre-Walker, A., & Keightley, P. D. (2007). The distribution of fitness effects of new mutations. Nature Reviews Genetics, 8(8), 610–618.

Glémin, S. (2007). Mating systems and the efficacy of selection at the molecular level. Genetics, 177(2), 905–916.

Goldberg, E. E., Kohn, J. R., Lande, R., Robertson, K. A., Smith, S. A., & Igić, B. (2010). Species selection maintains self-incompatibility. Science, 330(6003), 493–495.

Goodwillie, C., Kalisz, S., & Eckert, C. G. (2005). The evolutionary enigma of mixed mating systems in plants: occurrence, theoretical explanations, and empirical evidence. Annu. Rev. Ecol. Evol. Syst., 36, 47–79.

Haldane, J. B. (1919). The combination of linkage values and the calculation of distances between the loci of linked factors. J Genet, 8(29), 299–309.

Hartfield, M., Bataillon, T., & Glémin, S. (2017). The evolutionary interplay between adaptation and selffertilization. Trends in Genetics, 33(6), 420–431.

Hill W. G., & Robertson. A. (1966) The effect of linkage on limits to artificial selection. Genetic Research, 8, 269–294.

Hojsgaard, D., & Hörandl, E. (2019). The rise of apomixis in natural plant populations. Frontiers in Plant Science, 10, 358.

Honnay, O., & Jacquemyn, H. (2008). A meta-analysis of the relation between mating system, growth form and genotypic diversity in clonal plant species. Evolutionary Ecology, 22, 299–312.

Hörandl, E. (2010). The evolution of self-fertility in apomictic plants. Sexual plant reproduction, 23(1), 73–86.

Innes, D. J., Fox, C. J., & Winsor, G. L. (2000). Avoiding the cost of males in obligately asexual Daphnia pulex (Leydig). Proceedings of the Royal Society of London. Series B: Biological Sciences, 267(1447), 991–997.

Jarne, P., & Charlesworth, D. (1993). The evolution of the selfing rate in functionally hermaphrodite plants and animals. Annual Review of Ecology and Systematics, 24(1), 441–466.

Keightley, P. D., & Otto, S. P. (2006). Interference among deleterious mutations favours sex and recombination in finite populations. Nature, 443(7107), 89–92.

Klekowski, E. J., & Godfrey, P. J. (1989). Ageing and mutation in plants. Nature, 340(6232), 389–391.

Kondrashov, A. S. (1985). Deleterious mutations as an evolutionary factor. II. Facultative apomixis and selfing. Genetics, 111(3), 635–653.

Kron, P., Stewart, S. C., & Back, A. (1993). Self-compatibility, autonomous self-pollination, and insectmediated pollination in the clonal species Iris versicolor. Canadian Journal of Botany, 71(11), 1503–1509.

Kumpulainen, T., Grapputo, A., & Mappes, J. (2004). Parasites and sexual reproduction in psychid moths. Evolution, 58(7), 1511–1520.

Lande, R., & Schemske, D. W. (1985). The evolution of self-fertilization and inbreeding depression in plants. I. Genetic models. Evolution, 39(1), 24–40.

Lehtonen, J., Jennions, M. D., & Kokko, H. (2012). The many costs of sex. Trends in ecology & evolution, 27(3), 172–178.

Lewis W. M. Jr. 1987. The cost of sex. Pages 33–57 in The Evolution of Sex and its Consequences, edited by S. C. Stearns. Basel (Switzerland): Birkhäuser Verlag.

Marriage, T. N., & Orive, M. E. (2012). Mutation-selection balance and mixed mating with asexual reproduction. Journal of theoretical biology, 308, 25–35.

Mazer SJ, Delesalle VA, Paz H. 2007. Evolution of mating system and the genetic covariance between male and female investment in Clarkia (onagraceae): selfing opposes the evolution of trade-offs. Evolution 61: 83–98.

Meirmans, S., Meirmans, P. G., & Kirkendall, L. R. (2012). The costs of sex: facing real-world complexities. The Quarterly Review of Biology, 87(1), 19–40.

Michaels, H. J., & Bazzaz, F. A. (1986). Resource allocation and demography of sexual and apomictic Antennaria parlinii. Ecology, 67(1), 27–36.

Mione, T., & Anderson, G. J. (1992). Pollen-ovule ratios and breeding system evolution in Solanum section Basarthrum (Solanaceae). American Journal of Botany, 79(3), 279–287.

Muirhead, C. A., & Lande, R. (1997). Inbreeding depression under joint selfing, outcrossing, and asexuality. Evolution, 51(5), 1409–1415.

Mukai, T., Chigusa, S. I., Mettler, L. E., & Crow, J. F. (1972). Mutation rate and dominance of genes affecting viability in Drosophila melanogaster. Genetics, 72(2), 335–355.

Nakayama, Y., Seno, H., & Matsuda, H. (2002). A population dynamic model for facultative agamosperms. Journal of theoretical biology, 215(2), 253–262.

Nordborg, M., Charlesworth, B., & Charlesworth, D. (1996). The effect of recombination on background selection. Genetics Research, 67(2), 159–174.

Otto, S. P. (2003). The advantages of segregation and the evolution of sex. Genetics, 164(3), 1099–1118.

Otto, S. P. (2009). The evolutionary enigma of sex. The American Naturalist, 174(S1), S1–S14.

Otto, S. P., & Barton, N. H. (2001). Selection for recombination in small populations. Evolution, 55(10), 1921–1931.

Otto, S. P., & Payseur, B. A. (2019). Crossover interference: shedding light on the evolution of recombination. Annual review of genetics, 53, 19–44.

Richards, A. J. (2003). Apomixis in flowering plants: an overview. Philosophical Transactions of the Royal Society of London. Series B: Biological Sciences, 358(1434), 1085–1093.

Ritland, C., & Ritland, K. (1989). Variation of sex allocation among eight taxa of the Mimulus guttatus species complex (Scrophulariaceae). American Journal of Botany, 76(12), 1731–1739.

Roze, D. (2014). Selection for sex in finite populations. Journal of evolutionary biology, 27(7), 1304–1322.

Roze, D. (2021). A simple expression for the strength of selection on recombination generated by interference among mutations. Proceedings of the National Academy of Sciences, 118(19).

Roze, D., & Barton, N. H. (2006). The Hill–Robertson effect and the evolution of recombination. Genetics, 173(3), 1793–1811.

Roze, D., & Lenormand, T. (2005). Self-fertilization and the evolution of recombination. Genetics, 170(2), 841–857.

Roze, D., & Michod, R. E. (2010). Deleterious mutations and selection for sex in finite diploid populations. Genetics, 184(4), 1095–1112.

Sicard, A., & Lenhard, M. (2011). The selfing syndrome: a model for studying the genetic and evolutionary basis of morphological adaptation in plants. Annals of botany, 107(9), 1433–1443.

Silvertown, J. (2008). The evolutionary maintenance of sexual reproduction: evidence from the ecological distribution of asexual reproduction in clonal plants. International journal of plant sciences, 169(1), 157–168.

Simmons, M. J., & Crow, J. F. (1977). Mutations affecting fitness in Drosophila populations. Annual review of genetics, 11(1), 49–78.

Smith, J. M., & Maynard-Smith, J. (1978). The evolution of sex (Vol. 4). Cambridge: Cambridge University Press.

Soltis, D. E., & Soltis, P. S. (1992). The distribution of selfing rates in homosporous ferns. American Journal of Botany, 79(1), 97–100.

Stephan, W. (2010). Genetic hitchhiking versus background selection: the controversy and its implications. Philosophical Transactions of the Royal Society B: Biological Sciences, 365(1544), 1245–1253.

Stetsenko, R., & Roze, D. (2022). The evolution of recombination in self-fertilizing organisms. Genetics, 222(1), iyac114.

Sutherland, S. (1986). Patterns of fruit-set: What controls fruit-flower ratios in plants?. Evolution, 40(1), 117–128.

Uyenoyama, M. K., & Bengtsson, B. O. (1989). On the origin of meiotic reproduction: a genetic modifier model. Genetics, 123(4), 873–885.

Vallejo-Marín, M., Dorken, M. E., & Barrett, S. C. (2010). The ecological and evolutionary consequences of clonality for plant mating. Annual Review of Ecology, Evolution, and Systematics, 41, 193–213.

Vitalis, R., & Couvet, D. (2001). Two-locus identity probabilities and identity disequilibrium in a partially selfing subdivided population. Genetics Research, 77(1), 67–81.

Whitton, J., Sears, C. J., Baack, E. J., & Otto, S. P. (2008). The dynamic nature of apomixis in the angiosperms. International Journal of Plant Sciences, 169(1), 169–182.

Williams, G. C. (1975). Sex and evolution. Princeton Univ. Press, Princeton, NJ.

Wolinska, J., & Lively, C. M. (2008). The cost of males in Daphnia pulex. Oikos, 117(11), 1637–1646.

Xu, K. (2022). The genetic basis of selfing rate evolution. Evolution, 76(5), 883–898.

