## Supplementary Figures and Tables for "Effects of selfing on the evolution of sexual reproduction"

**Supplementary Figures for "Effects of selfing on the evolution of sexual reproduction"**

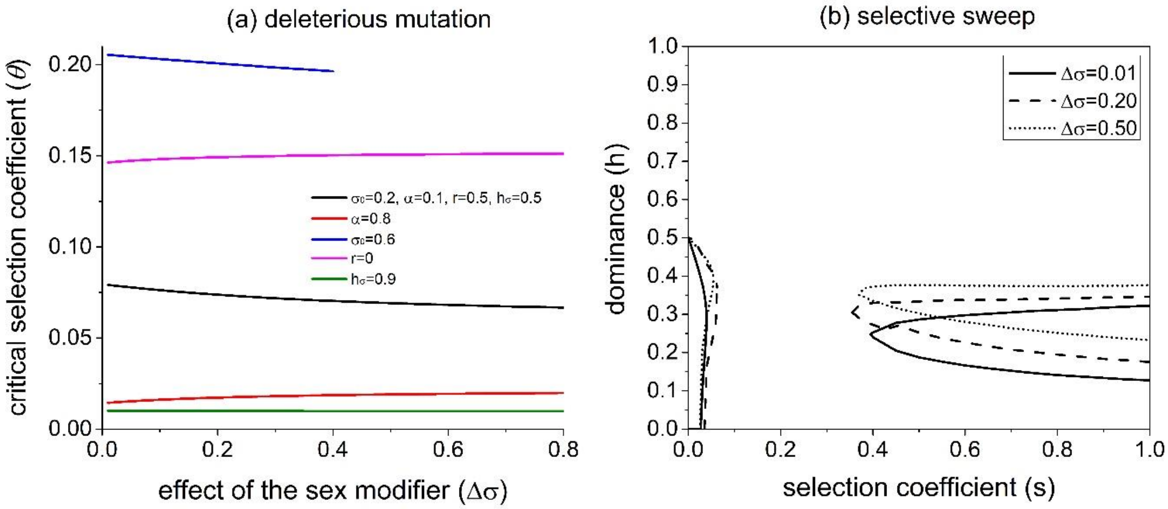


**Figure S1**. The influences of the effect of a sex modifier ($\Delta\sigma$) on its invasion conditions. Panel (a) shows the effects of $\Delta\sigma$ on the critical selection coefficient $\theta$ at which the modifier cannot invade for any dominance $h$ for the case when the fitness locus is at selection-mutation balance ($\theta$ is given in Section 3.3 of the Mathematica Notebook). The black line is taken as the standard parameter values, and other lines differ from it by one of the parameters. Panel (b) shows the case when the fitness locus experiences selective sweep, and $p_{A,0} = 0.001, p_{A,T} = 0.999, p_{m,0} = 0.999$. Other parameters are $\mu = 10^{-5}, c = 1, \tau = 0$.

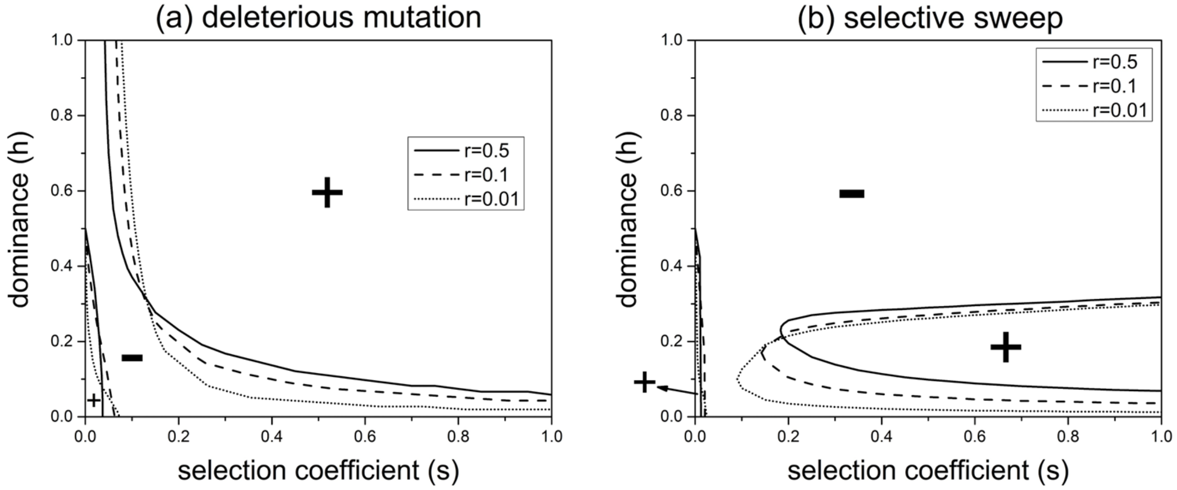


**Figure S2.** Effects of the recombination rate ($r$) on the evolution of a modifier that increases the sexual reproduction rate ($\Delta\sigma = 0.01$, $h_\sigma = 0.05$) when allele $a$ at the fitness locus is deleterious (panel (a)) or when the fitness locus experiences selective sweep (panel (b)). The meanings of "+" and "-" are the same as in Figure 1. Other parameters are $\sigma_0 = 0.1, \alpha = 0.1, \mu = 10^{-5}, c = 1, \tau = 0, p_{A,0} = 0.001, p_{A,T} = 0.999, p_{m,0} = 0.999$.

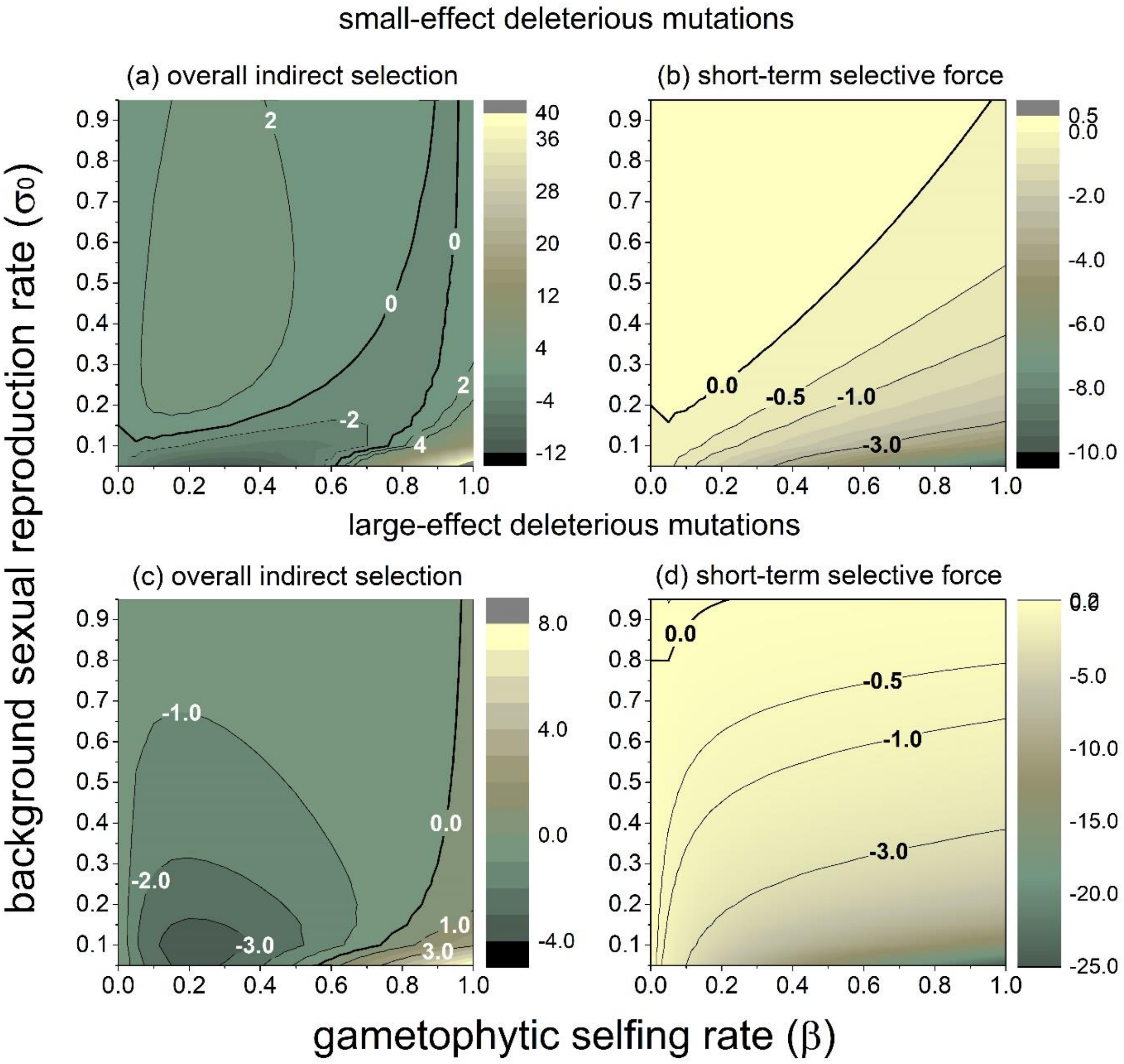


**Figure S3.** Strength of indirect selection generated by deleterious mutations in the genome due to intra-locus interactions on a modifier increasing the rate of sexual reproduction under gametophytic selfing. The upper and bottom panels show selection caused by small-effect (upper panels) and large-effect (bottom panels) deleterious mutations. Panels (a) and (c) show the overall strength of indirect selection. Panels (b) and (d) show the strength of short-term (dis)advantage of the modifier due to fitness differences between sexually and asexually reproduced offspring. Parameters used are the same as Figure 3.

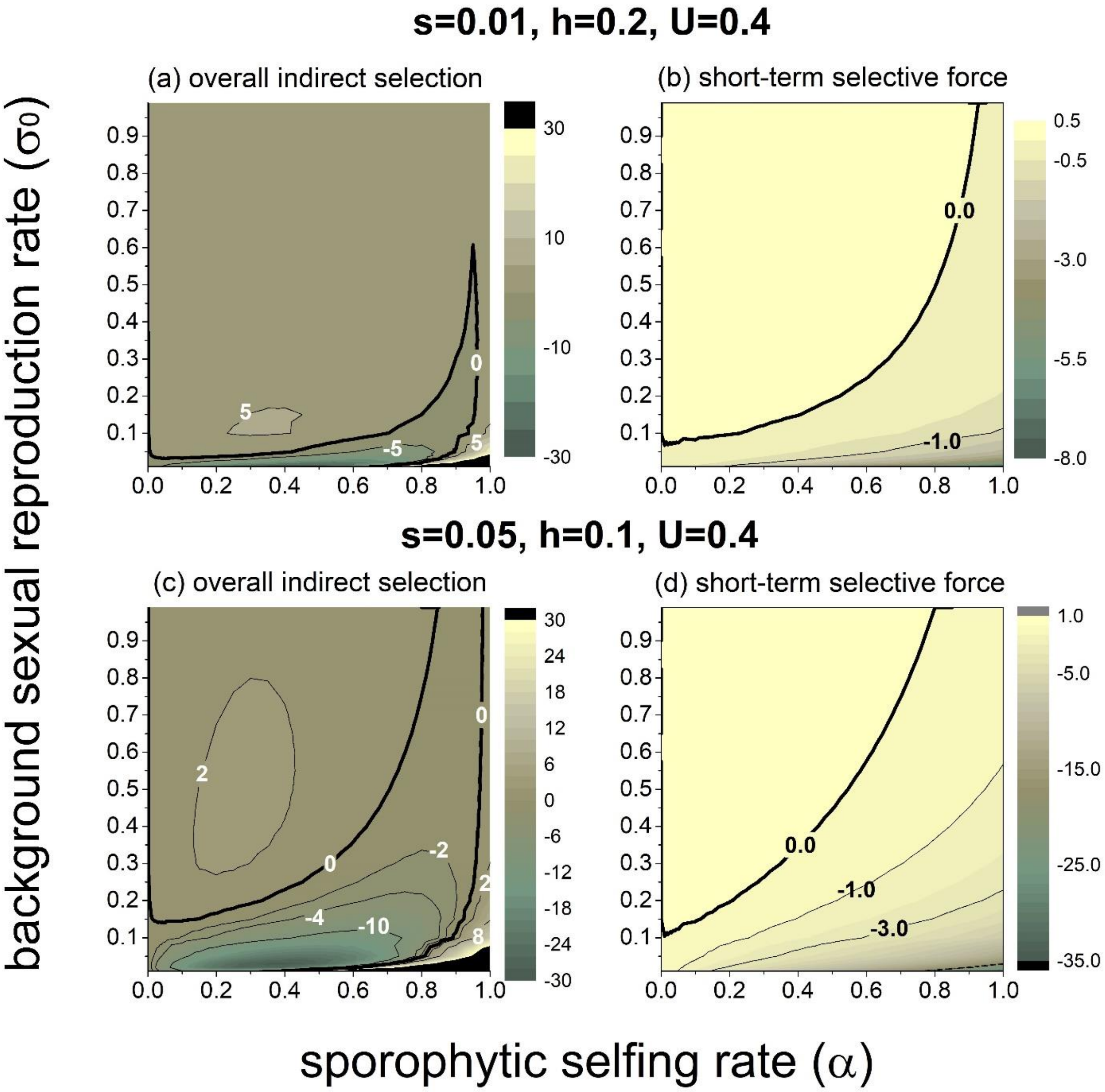


**Figure S4**. A modifier increasing the rate of sexual reproduction is favored in a larger parameter space when deleterious mutations have smaller effect (compare Fig. S4(a) vs. 3(a)) and more recessive (compare Fig. S4(c) with 3(c)). See the caption of Fig. 3 for interpretations. $s = 0.01$ for Other parameters are the same as Fig. 3.
