## Supplementary Materials for "Effects of selfing on the evolution of sexual reproduction"

Here I show how the sexual reproduction rate and the selfing rate affect the relative fitness of asexually reproduced and selfed offspring under sporophytic and gametophytic selfing, as well as the mean fitness at a fitness locus.

### I. Sporophytic selfing

To see how the sexual reproduction rate $\sigma_0$ and selfing rate $\alpha$ affect the relative fitness reduction of asexually reproduced offspring $d_a$, based on equation (12a), we have

$$\frac{\partial d_a}{\partial \sigma} = \frac{Fhs(1+F)}{\big((1+F-\sigma_0)hs + (F+h-hF)\sigma_0\big)^2}(1-2h)\mu, \tag{A1a}$$

$$\frac{\partial d_a}{\partial \alpha} = \frac{(1+F-Fs)^2 h\sigma_0}{2(F+h-Fh)^2(s(1-\sigma_0)+\sigma_0)}(1-2h)\mu. \tag{A1b}$$

When $h < 0.5$, both equations are greater than 0. Therefore, the relative fitness of asexually reproduced offspring is lower when $\sigma_0$ and $\alpha$ are higher. Similarly, for the inbreeding depression of selfed offspring given in equation (12b), we have

$$\frac{\partial d_s}{\partial \sigma} = -\frac{F(1+F)(1-2h)^2 s}{2\big((1+F-\sigma_0)hs + (F+h-hF)\sigma_0\big)^2}\mu < 0, \tag{A2a}$$

$$\frac{\partial d_s}{\partial \alpha} = -\frac{(1-2h)^2(1+F-Fs)^2\sigma_0}{4(F+h-Fh)^2(s(1-\sigma_0)+\sigma_0)}\mu < 0. \tag{A2b}$$

Therefore, the relative fitness of selfed offspring is higher when the rate of sexual reproduction and the selfing rate are higher.

Finally, I analyze the effects of selfing rate $\alpha$ and the background sexual reproduction rate $\sigma_0$ on the mean fitness of the population, which is given by

$$\overline{w}(\sigma_0,\alpha) = \hat{x}_{11} + \hat{x}_{12}(1-hs) + \hat{x}_{22}(1-s) = 1 - \frac{F+2h(1-F)}{h+(1-h)F}\mu, (\alpha > 0), \tag{A3}$$

Note that the above approximation only applies to the case when $\alpha > 0$. Under outcrossing ($\alpha = 0$), $\overline{w}$ should be expanded to the second order $\mu^2$. To see how sexual reproduction rate affects the mean fitness, and how selfing affects the strength of selection on sexual reproduction caused by between-population selection, note that

$$\frac{\partial \overline{w}(\sigma_0, \alpha)}{\partial \sigma_0} = \frac{Fh\big(1+F(1-s)\big)s\mu}{(F+h-Fh)^2(s(1-\sigma_0)+\sigma_0)\sigma_0}\mu > 0 \tag{A4a}$$

$$\frac{\partial^2 \overline{w}(\sigma_0, \alpha)}{\partial \sigma_0 \partial \alpha} = \frac{hs(1+F-Fs)^2(Fh(3-2s)+h-F)\mu}{2(s+\sigma_0-s\sigma_0)^2\big(h+F(1-h)\big)^3} \tag{A4b}$$

Equation (4a) shows that a higher sexual reproduction rate $\sigma_0$ will increase the mean fitness, so sexual reproduction is favored by between-population selection. Equation (4b) shows that selfing strengthens this selection when $h > \frac{F}{1+F(3-2s)}$. Considering deleterious mutations in the genome, for small-effect deleterious mutations, since they are usually partially recessive, selfing tends to increase the selective strength on sexual reproduction. In contrast, for large-effect lethal mutations, since they are often highly recessive, selfing is likely to reduce selective strength. Nevertheless, since the genomic mutation rate of large-effect lethal allele is much smaller than small-effect mutations, the genetic load is mainly contributed by small-effect mutations in the genome. Therefore, generally, selfing may increase the strength of selection on sexual reproduction caused by selection between populations.

**II. Gametophytic selfing**

Under gametophytic selfing, the fitness reduction of asexually reproduced and selfed offspring relative to outcrossed offspring caused by a single locus is respectively

$$d_a = \frac{(1-2h)\beta\sigma}{hs+\sigma\beta+(1-s-\beta)h\sigma}\mu, \tag{A5a}$$

$$d_s = \frac{(1-2h)(s(1-\sigma)+\sigma)}{hs+\sigma\beta+(1-s-\beta)h\sigma}\mu. \tag{A5b}$$

The fitness reduction of selfed offspring relative to asexually reproduced offspring is

$$\frac{d_s}{d_a} = \frac{1}{\beta}\left(1+\frac{1-\sigma}{\sigma}s\right) \geq 1. \tag{A6}$$

Therefore, selfed offspring have lower fitness than asexually reproduced offspring.

Taking the first order derivative of $d_a$ and $d_s$ for $\sigma$ and $\beta$, and $h < 0.5$, we have

$$\frac{\partial d_a}{\partial \sigma} = \frac{\beta s h(1-2h)}{(\beta\sigma + h(s+\sigma-(s+\beta)\sigma))^2}\mu > 0, \tag{A7a}$$

$$\frac{\partial d_a}{\partial \beta} = \frac{h(1-2h)(s(1-\sigma)+\sigma)\sigma}{(\beta\sigma + h(s+\sigma-(s+\beta)\sigma))^2}\mu > 0, \tag{A7b}$$

and

$$\frac{\partial d_s}{\partial \sigma} = -\frac{(1-h)(1-2h)s\alpha}{\big(\beta\sigma + h(s+\sigma-(s+\beta)\sigma)\big)^2}\mu < 0, \tag{A8a}$$

$$\frac{\partial d_s}{\partial \beta} = -\frac{(1-h)(1-2h)(s(1-\sigma)+\sigma)\sigma}{\big(\beta\sigma + h(s+\sigma-(s+\alpha)\sigma)\big)^2}\mu < 0. \tag{A8b}$$

Therefore, as the rate of sexual reproduction and the gametophytic selfing rate increases, the relative fitness of asexually reproduced offspring decreases, while the relative fitness of selfed offspring increases. Also, it can be shown that the genetic load of sexually reproduced offspring relative to asexually reproduced offspring is

$$\frac{1-w_s}{1-w_a} \approx \beta\frac{d_s}{d_a} = \begin{cases} 1, & \beta = 0 \\ 1 + s\left(\dfrac{1}{\sigma} - 1\right) \geq 1, & \beta > 0 \end{cases}. \tag{A9}$$

Equation (A9) shows that similar to sporophytic selfing, when deleterious mutations are recessive, the presence of gametophytic selfing causes sexually reproduced offspring to be less fit than asexually reproduced offspring due to increased homozygosity. This is because sexually reproduced offspring always have higher homozygosity than asexually reproduced offspring, since the inbreeding coefficient within asexually and sexually reproduced offspring is respectively, $\frac{\sigma}{\sigma+(1-\sigma)s}\beta$ and $\beta$. However, equation (A9) shows that the fitness of sexually reproduced offspring relative to asexually reproduced offspring is independent of the gametophytic selfing rate $\beta$. In contrast, under sporophytic selfing, a higher selfing rate will reduce the relative fitness of sexually reproduced offspring (see equation (12)).
